# *InMYB21B* Promotes Petal Cell Expansion and Flower Opening in Japanese Morning Glory (*Ipomoea nil*)

**DOI:** 10.64898/2026.08.22.746480

**Authors:** Soya Nakagawa, Atsushi Hoshino

## Abstract

Flower opening is a complex developmental process involving coordinated changes in cell proliferation and cell expansion. Although several regulators of flower opening have been identified, how transcriptional programs are coordinated with the cellular and metabolic changes underlying petal expansion immediately before flower opening remains incompletely understood. Japanese morning glory (*Ipomoea nil*) is a suitable model for investigating these processes because its flowers open synchronously at a predictable time. This study aimed to identify transcriptional regulators involved in petal development and flower opening in Japanese morning glory. Temporal analyses of petal growth, sugar metabolism, and gene expression revealed that petal development was driven by both cell proliferation and cell expansion until approximately 48 h before flower opening, whereas cell expansion predominated thereafter. Weighted gene co-expression network analysis identified two genes encoding R2R3-MYB subgroup 19 transcription factors, *InMYB21A* and *InMYB21B*, as candidate regulators associated with petal development. CRISPR/Cas9-mediated knockout analysis revealed a prominent role for *InMYB21B*, whose loss markedly impaired petal cell expansion and prevented flower opening. *InMYB21B* knockout also impaired stamen and pistil development, resulting in male and female sterility. Starch degradation and glucose accumulation were impaired in *InMYB21B* knockout petals. Transcriptome analysis revealed delayed transcriptomic progression during petal development and reduced expression of genes associated with starch degradation, sucrose metabolism, cell wall remodeling, and water transport. These findings identify *InMYB21B* as a key regulator of petal cell expansion and flower opening in Japanese morning glory and show that loss of *InMYB21B* disrupts both metabolic and transcriptomic progression during late petal development.

## INTRODUCTION

Seed plants develop floral organs for reproduction. In self-pollinating plants, bisexual flowers bearing both male and female reproductive organs are formed, and the development of petals, pistils, and stamens proceeds in a coordinated manner to ensure successful self-pollination. Floral organ formation proceeds through the following three stages. In the first stage, the identity of each floral organ is established. Genetic studies using *Arabidopsis thaliana* and *Antirrhinum majus* as model plants have identified genes encoding MADS-box transcription factors belonging to the ABC classes as key regulators of this process (Bowman and Moyroud 2024; Immink et al. 2010). In the second stage, cell division becomes active, resulting in an increase in cell number within each organ. Numerous genes involved in the cell cycle and cell division have been identified in plants, and studies using floral organ size mutants have revealed genes that regulate the activity and duration of cell division, particularly in petals (Hepworth and Lenhard 2014; Irish 2008; Krizek and Anderson 2013). In the third stage, cell division activity decreases, while cell expansion becomes predominant (Guan et al. 2025; Wang et al. 2024). Cell expansion is thought to be driven by water uptake associated with increased intracellular osmotic pressure and by cell wall remodeling (Johnson and Lenhard 2011; van Doorn and Van Meeteren 2003).

The increase in osmotic pressure in petal cells is caused by the metabolism of storage polysaccharides, such as starch and fructans, and sucrose transported to the petals through the phloem (van Doorn and Kamdee 2014; van Doorn and Van Meeteren 2003). In many plant species, starch accumulation in petal cells has been reported, and in lily, inhibition of starch degradation suppresses petal development (Bieleski et al. 2000). Furthermore, sucrose transported to petals is hydrolyzed into glucose and fructose by the irreversible reaction catalyzed by invertase (INV), or converted into fructose and UDP-glucose through the reversible reaction catalyzed by sucrose synthase (SUS) (Stein and Granot 2019; Wan et al. 2018). In particular, vacuolar INV activity and expression have been associated with flower development and opening in rose and lisianthus (Farci et al. 2016; Harada et al. 2021). SUS has also been implicated in flower development in cucumber (Fan et al. 2019), whereas flower opening in azalea has been associated with source–sink sucrose metabolism (Christiaens et al. 2016). Cell wall remodeling is another important process that promotes increases in cell size associated with increased turgor pressure. Major enzymes involved in this process include xyloglucan endotransglucosylase/hydrolase (XTH) and expansin (EXP). XTH cleaves xyloglucan chains, a component of the cell wall, thereby promoting cell wall loosening (Eklof and Brumer 2010). In contrast, EXP promotes cell wall loosening by disrupting hydrogen bonds between cell wall polysaccharides in a pH-dependent manner (Marowa et al. 2016; Sampedro and Cosgrove 2005). In particular, the expression of XTH and EXP genes has been associated with petal growth and flower opening in gerbera, Japanese morning glory, four-o’clock, and carnation (Gookin et al. 2003; Harada et al. 2011; Laitinen et al. 2007; Shinozaki et al. 2014).

Subgroup 19 (SG19) of the R2R3-MYB transcription factor family has diverse functions in floral organ development, with roles that have diversified among plant species. Previous studies have characterized the functions of SG19 transcription factors mainly in Arabidopsis, rice, petunia, and tomato, demonstrating their important roles in flower opening (Chopy et al. 2023; Colquhoun et al. 2011; Gou et al. 2024; Liu and Thornburg 2012; Niwa et al. 2018; Reeves et al. 2012; Schubert et al. 2019), senescence (Chopy et al. 2023; Colquhoun et al. 2011), stamen development (Colquhoun et al. 2011; Mandaokar et al. 2006; Niwa et al. 2018; Reeves et al. 2012), anther dehiscence (Mandaokar et al. 2006; Song et al. 2011), pollen development (Mandaokar et al. 2006; Niwa et al. 2018; Song et al. 2011), nectary development (Chopy et al. 2023; Liu and Thornburg 2012; Reeves et al. 2012), ovule development (Schubert et al. 2019), pistil development (Chopy et al. 2023; Colquhoun et al. 2011; Niwa et al. 2018; Reeves et al. 2012; Yarahmadov et al. 2020), and fragrance production (Colquhoun et al. 2011; Reeves et al. 2012; Spitzer-Rimon et al. 2010). *EOB2*, which encodes an SG19 transcription factor in petunia (*Petunia axillaris*), is involved in petal maturation, with loss of its transcriptional activation domain resulting in reduced petal cell size and impaired starch degradation (Chopy et al. 2023). In rice, *OsMYB8*, which encodes an SG19 transcription factor, promotes floret opening by directly activating *OsJAR1*, thereby affecting JA-Ile accumulation and the expression of genes associated with lodicule hydration and cell wall remodeling (Gou et al. 2024).

R2R3-MYB SG19 transcription factors contain two major functional domains: an N-terminal R2R3-MYB domain and a C-terminal transcriptional activation domain (Chopy et al. 2023; Liu et al. 2009; Schubert et al. 2019). The R2R3-MYB domain is known to have three major functions. First, it mediates interactions with proteins such as JAZ, DELLA, and bHLH proteins (Huang et al. 2017; Huang et al. 2020; Qi et al. 2015; Schubert et al. 2019; Song et al. 2011; Yang et al. 2020). In Arabidopsis, JAZ and DELLA proteins negatively regulate the activity of MYB SG19 transcription factors through interactions with the R2R3-MYB domain (Huang et al. 2017; Huang et al. 2020; Song et al. 2011). In contrast, MYC2, MYC3, MYC4, and MYC5, which belong to subgroup IIIe of the bHLH transcription factor family, form bHLH–MYB complexes with MYB SG19 transcription factors and regulate the transcription of target genes together with these MYB proteins (Qi et al. 2015; Yang et al. 2020). Second, the R2R3-MYB domain of SG19 transcription factors has been reported to recognize cis-regulatory sequences related to [G/A]TT[A/T]GG[T/C] (Medina-Puche et al. 2015; Weirauch et al. 2014). Recognition of similar cis-regulatory sequences by R2R3-MYB SG19 transcription factors has been reported in diverse plant species, including Arabidopsis, rice, Japanese apricot, strawberry, and freesia (Gou et al. 2024; Medina-Puche et al. 2015; Weirauch et al. 2014; Yang et al. 2020; Yuan et al. 2024). Third, the nuclear localization signal (NLS) within the R2R3-MYB domain mediates nuclear localization of the translated MYB SG19 transcription factors (Liu et al. 2009). The C-terminal transcriptional activation domain functions to recruit the transcriptional machinery and is broadly conserved from Solanaceae to Brassicaceae (Chopy et al. 2023).

Japanese morning glory (*Ipomoea nil*) is a valuable biological resource that has long been used as a model for physiological studies of flower opening. Its whole-genome sequence has been published (Hoshino et al. 2016), and an efficient genome-editing method has also been established (Watanabe et al. 2017). Flower opening in Japanese morning glory is controlled by the circadian clock, with flowers opening synchronously in the morning and wilting on the same day. In addition, from approximately four days before flower opening, the time remaining until flower opening can be readily estimated from petal size. Taking advantage of these characteristics, time-series transcriptome analysis was performed on petals at 3-h intervals from 72 h before flower opening to 12 h after flower opening (Nakagawa et al. 2025). In the present study, this time-series transcriptome dataset was used to perform weighted gene co-expression network analysis (WGCNA) to elucidate the gene networks underlying petal development in Japanese morning glory. *InMYB21A* and *InMYB21B*, which encode R2R3-MYB SG19 transcription factors, were identified as hub genes. Functional analysis using CRISPR/Cas9-mediated genome editing demonstrated that *InMYB21B* plays a central role in petal cell expansion and flower opening, as well as an important role in pistil and stamen development.

## RESULTS

### Sugar metabolism and cell wall remodeling are associated with petal expansion in Japanese morning glory

Petal fresh weight and length were measured at 6-h intervals from 69 h before flower opening to 9 h after flower opening (Figure 1A). Petal fresh weight and length gradually increased from 69 to 21 h before flower opening and then increased rapidly from 21 to 3 h before flower opening. To investigate the contribution of sugar metabolism to petal expansion, the contents of glucose, sucrose, and starch were determined using enzymatic assays (Figure 1B). Starch content decreased from 21 to 3 h before flower opening, whereas glucose content increased from 27 to 15 h before flower opening. Sucrose, a major translocated sugar, increased from 27 to 21 h before flower opening. These results suggest that the rapid expansion of petals beginning approximately 21 h before flower opening is associated with starch degradation and the accumulation of glucose and sucrose, which may contribute to increased intracellular osmotic pressure and petal cell expansion.

**Figure 1.**
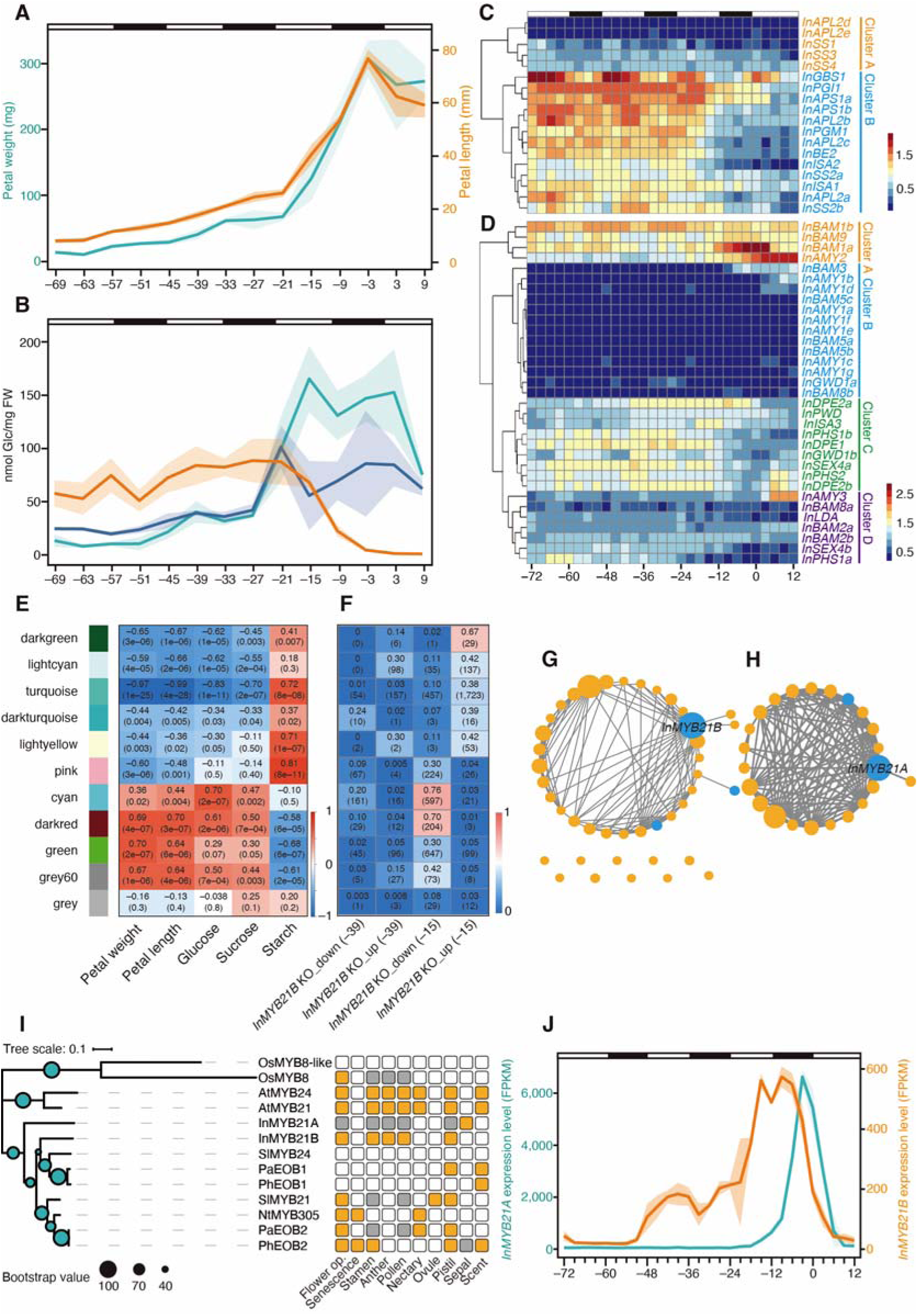
Identification of *InMYB21A* and *InMYB21B* as petal development-associated genes (A) Temporal changes in petal fresh weight (green) and length (orange). The x-axis indicates time relative to flower opening (0 h). The bars at the top indicate light conditions, with white and black representing light and dark periods, respectively. Mean values at each time point are shown as lines, and standard deviations are indicated by shaded areas (±SD; n = 3). (B) Temporal changes in glucose (green), sucrose (blue), and starch (orange) contents determined by enzymatic assays. The x-axis indicates time relative to flower opening (0 h), and the y-axis indicates glucose equivalents (nmol Glc/mg FW). The bars at the top indicate light conditions, with white and black representing light and dark periods, respectively. Mean values at each time point are shown as lines, and standard deviations are indicated by shaded areas (±SD; n = 3). Heatmaps showing the temporal expression patterns of starch biosynthesis genes (C) and starch degradation genes (D). The gene list is provided in Table S1. Expression levels are shown as log10(FPKM + 1). The x-axis indicates time relative to flower opening (0 h). The bars at the top indicate light conditions, with white and black representing light and dark periods, respectively. Gene names are shown in different colors according to their respective clusters. (E) Results of weighted gene co-expression network analysis (WGCNA). Correlations between each co-expression module and physiological traits, including petal fresh weight, length, glucose content, sucrose content, and starch content. Values within the cells indicate correlation coefficients, with *p* values shown in parentheses. The color of each cell corresponds to the color scale indicating the magnitude of the correlation coefficient. (F) Proportion of genes in each co-expression module identified as differentially expressed genes (DEGs). The four columns, from left to right, represent downregulated DEGs at 39 h, upregulated DEGs at 39 h, downregulated DEGs at 15 h, and upregulated DEGs at 15 h before flower opening in *InMYB21B* KO line #2. Values within the cells indicate the proportion of DEGs relative to the total number of genes in each co-expression module, with the number of genes shown in parentheses. The color of each cell corresponds to the color scale indicating the magnitude of the proportion. (G, H) Co-expression networks of transcriptional regulatory genes in the cyan (G) and darkred (H) co-expression modules. Blue and orange circles represent nodes corresponding to floral organ-specific and non-specific genes, respectively. Gray lines represent co-expression relationships. Larger nodes indicate a greater number of co-expressed genes and are inferred to function as hub genes. (I) Phylogenetic analysis and reported functions of R2R3-MYB subgroup 19 transcription factors from *Oryza sativa* (Os), *Arabidopsis thaliana* (At), *Ipomoea nil* (In), *Solanum lycopersicum* (Sl), *Petunia axillaris* (Pa), *Petunia hybrida* (Ph), and *Nicotiana tabacum* (Nt). The sizes of the blue circles at the nodes indicate bootstrap values, and the scale bar indicates the number of amino acid substitutions per site. The color coding on the right indicates reported functions in each floral organ (orange, function reported; gray, no function reported; white, not reported). (J) Temporal expression patterns of *InMYB21A* and *InMYB21B*. The x-axis indicates time relative to flower opening (0 h). The left and right y-axes indicate FPKM values for *InMYB21A* and *InMYB21B*, respectively. Mean FPKM values at each time point are shown as lines, and standard deviations are indicated by shaded areas (±SD; n = 3). The bars at the top indicate light conditions, with white and black representing light and dark periods, respectively.

OrthoFinder analysis was used to identify Japanese morning glory orthologs of Arabidopsis genes involved in starch biosynthesis and degradation (Table S1). The expression patterns of these starch biosynthesis and degradation genes in petals were then analyzed using a time-series transcriptome dataset obtained from Japanese morning glory petals at 3-h intervals from 72 h before flower opening to 12 h after flower opening (Figures 1C and 1D). Starch biosynthesis genes were classified into two clusters (Clusters A and B) according to their temporal expression patterns. Genes in Cluster A showed relatively low expression levels, whereas those in Cluster B maintained high expression levels until 12 h before flower opening (Figure 1C). These results suggest that genes associated with starch biosynthesis remain highly expressed in Japanese morning glory petals until approximately 12 h before flower opening. In addition, *InGBS1*, *InAPS1a*, *InAPS1b*, and *InAPL2b* exhibited diurnal changes in expression, with higher expression levels during the light period (Figure 1C). Among the starch degradation genes, *InBAM1*a exhibited a transient increase in expression, peaking around flower opening (Figure 1D). *InBAM1*a showed high expression levels from 6 h before flower opening to 3 h after flower opening. Among the nine *BAM* genes in Arabidopsis, *BAM1* and *BAM3* are known to exhibit particularly high enzymatic activities (Streb and Zeeman 2012). Because *InBAM1a* is an ortholog of *BAM1*, it may contribute to starch degradation immediately before flower opening.

*INV* and *SUS* genes involved in sucrose metabolism form gene families in Arabidopsis, and their orthologs in Japanese morning glory were identified (Tables S2 and S3). Analysis of the temporal expression patterns of *INV* and *SUS* genes in petals revealed that two genes encoding vacuolar invertases, *InVIN1a* and *InVIN1d*, as well as *InSUS1b* and *InSUS1d*, were highly expressed in petals (Figure S1). Among these genes, *InVIN1d* and *InSUS1d* showed particularly high and transient increases in expression, with FPKM values exceeding 1,000 at 12 h before flower opening and at flower opening, respectively. Because vacuolar invertases are localized in the vacuole (Wan et al. 2018), whereas SUS proteins are localized in the cytoplasm (Stein and Granot 2019), these enzymes may contribute to sucrose metabolism in the vacuole and cytoplasm, respectively, immediately before flower opening, thereby increasing the osmotic pressure of petal cells. In contrast, sucrose is synthesized by sucrose-phosphate synthase (SPS) and sucrose-phosphate phosphatase (SPP) (Bagnato et al. 2023; Volkert et al. 2014). Orthologs of Arabidopsis genes encoding these enzymes were identified (Table S4), and analysis of their expression patterns revealed that *InSPSA1* and *InSPP1* were highly expressed at flower opening (Figure S1). These expression patterns suggest that *de novo* sucrose biosynthesis may also occur in petals around flower opening.

The expression levels of four *XTH* genes (*InXTH4*, *InXTH6*, *InXTH15*, and *InXTH22a*) and three *EXP* genes (*InEXPA1f*, *InEXPA1h*, and *InEXPA3b*) in Japanese morning glory petals have previously been analyzed by RT-qPCR, and the *XTH* genes were reported to exhibit increased expression at flower opening (Shinozaki et al. 2014). Because only a subset of *XTH* and *EXP* genes had previously been analyzed in Japanese morning glory, their Arabidopsis orthologs were comprehensively identified, and their expression patterns were examined (Tables S5, S6, and S7). Among the 21 identified *XTH* orthologs, the expression patterns of the four genes examined in the previous study were generally consistent with those reported previously. In addition, *InXTH22b* and *InXTH33a* showed increased expression, peaking at 6 h before flower opening (Figure S2). In particular, *InXTH4*and *InXTH6* exhibited exceptionally high and transient expression, with FPKM values of 7,253 at 3 h before flower opening and 16,944 at flower opening, respectively, suggesting that these genes may contribute to petal expansion associated with flower opening. Analysis of the expression patterns of *EXP* genes revealed that *InEXPA1f*, which had previously been reported to be below the detection limit (Shinozaki et al. 2014), showed the highest expression level among the *EXP* genes analyzed, with a high peak of 2,510 FPKM at 15 h before flower opening. Taken together, these expression patterns suggest that *InBAM1a*, *InVIN1d*, *InSUS1d*, *InXTH4*, *InXTH6*, and *InEXPA1f* may contribute to the increase in osmotic pressure and cell wall remodeling associated with flower opening.

### *InMYB21B* is a hub gene associated with petal expansion

Weighted gene co-expression network analysis (WGCNA) was performed to identify genes associated with changes in petal fresh weight, petal length, and sugar content (Figure 1E). Based on their expression patterns, genes were hierarchically clustered and subsequently assigned to 11 merged co-expression modules (Figure S3). Among these modules, the cyan, darkred, green, and grey60 modules showed positive correlations with petal fresh weight, petal length, glucose content, and sucrose content, but negative correlations with starch content (Figure 1E).

To examine the temporal expression patterns of genes in each module, the genes were compared with six clusters identified by time-series K-means clustering based on their expression patterns (Figure S4; Table S8). Among the genes assigned to the cyan, darkred, and grey60 co-expression modules, 493 genes (62.6%), 183 genes (63.1%), and 95 genes (54.3%), respectively, were classified into Cluster 4 (Table S9). Genes in Cluster 4 exhibited peak expression around flower opening, a pattern expected to be associated with petal expansion. In contrast, 1,236 genes (57.6%) in the green co-expression module, which showed the strongest association with petal fresh weight, were classified into Cluster 1 (Table S9). Genes in Cluster 1 exhibited peak expression at 12 h after flower opening.

Next, transcriptional regulatory genes within these four co-expression modules were predicted using the PlantTFDB database (Jin et al. 2017) (Table S10). Transcriptome data from six tissues—embryos, floral organs, leaves, roots, seed coats, and stems—reported by Hoshino et al. (2016) were used to further identify genes specifically expressed in floral organs, defined as genes with TPM values > 10 in floral organs and < 1 in each of the other five tissues (Table S10). Two floral organ-specific transcriptional regulatory genes were identified in the cyan module and one in the darkred module, whereas none were identified in the green or grey60 modules (Table S10). Focusing on the cyan and darkred co-expression modules, co-expression networks were constructed using only transcriptional regulatory genes (Figures 1G and 1H). In the cyan co-expression network, *InMYB21B* (INIL05g32166) was identified as a hub gene, whereas *InMYB21A* (INIL11g09839) was identified as a hub gene in the darkred co-expression network.

Maximum-likelihood phylogenetic analysis revealed that InMYB21A and InMYB21B belong to the R2R3-MYB SG19 transcription factor group (Figure 1I). An 11-amino-acid transcriptional activation domain at the C terminus of R2R3-MYB SG19 transcription factors is broadly conserved from Brassicaceae to Solanales (Chopy et al. 2023; Liu et al. 2009). However, InMYB21A contains a single serine insertion within this region (Figure S5A), which may impair its transcriptional activation activity.

The temporal expression patterns of *InMYB21A* and *InMYB21B* were also examined using the time-series transcriptome dataset obtained from Japanese morning glory petals (Figure 1J). *InMYB21A* exhibited a transient increase in expression, with a peak at 3 h before flower opening. In contrast, *InMYB21B* expression began to increase approximately 51 h before flower opening and reached its peak expression at 6 h before flower opening. Both genes were expressed in the limb, ray, tube, and stigma, whereas *InMYB21B* was additionally expressed in anthers (Figure S5B). Comparison of the promoter sequences of *InMYB21A* and *InMYB21B* using Harr plots revealed no apparent sequence similarity, providing no clear explanation for their distinct expression patterns (Figure S5C).

### Loss of *InMYB21B* impairs petal, pistil, and stamen development

To investigate the functions of *InMYB21A* and *InMYB21B*, loss-of-function knockout (KO) lines were generated using CRISPR/Cas9-mediated genome editing (Figure 2A; Figures S6–S9). For *InMYB21A* KO lines, individuals carrying the induced mutations were selected and self-pollinated, after which null segregants lacking the introduced transgene were selected (Figures S6 and S8). In contrast, *InMYB21B* KO lines exhibited sterility in the genome-edited T0 generation. Reciprocal crosses between five *InMYB21B* KO lines and wild-type plants were attempted, but no seeds were obtained, indicating that *InMYB21B* KO lines exhibited both male and female sterility. Therefore, biallelic mutant individuals carrying mutations in the regions encoding the MYB domains on both alleles were selected in the T0 generation (Figures S7 and S8).

**Figure 2.**
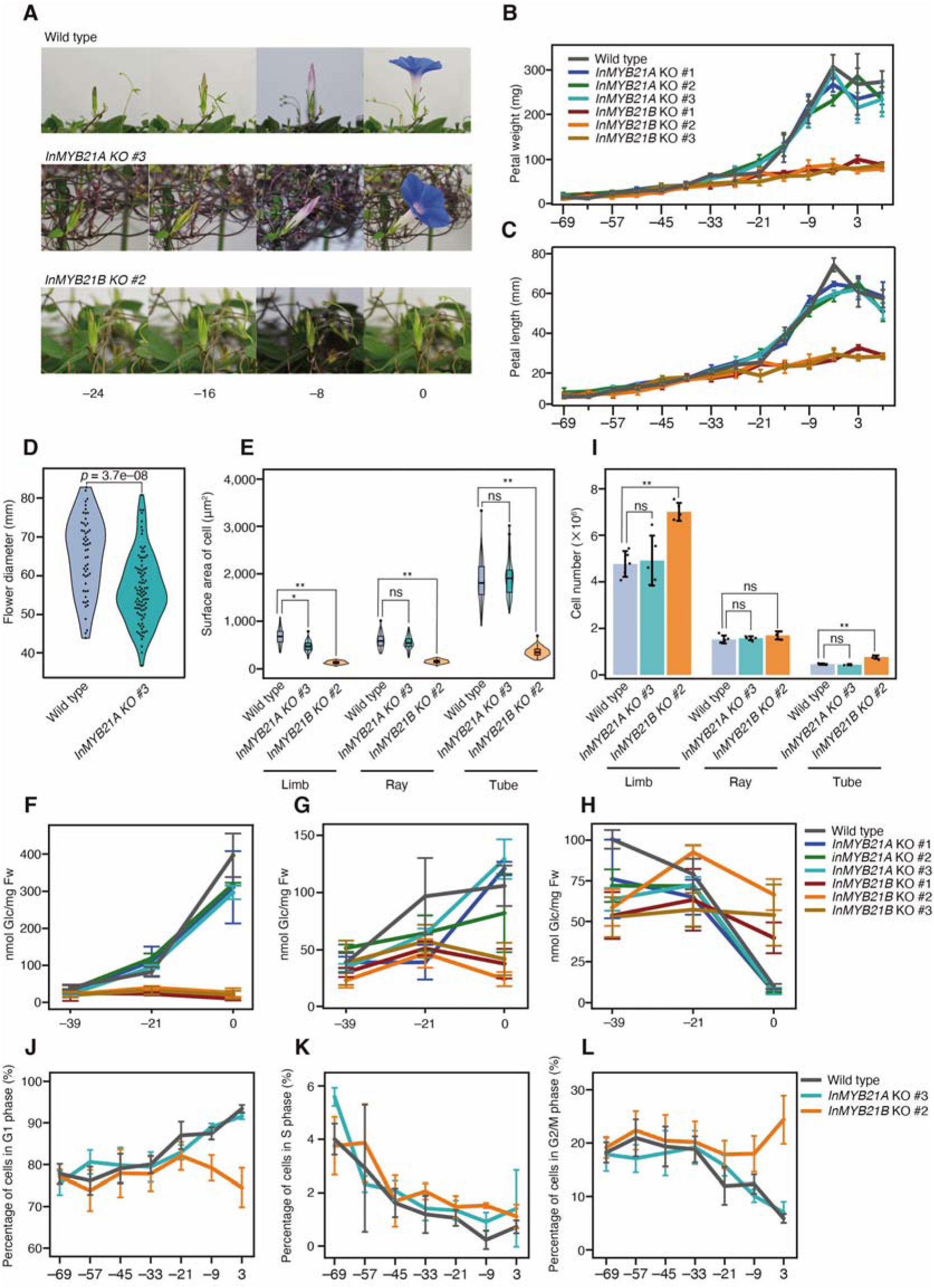
Functional analysi s of InMYB21A and InMYB21B by CRISPR/Cas9-mediated genome editing (A) Time-lapse images showing flower opening in wild-type plants, *InMYB21A* KO line #3, and *InMYB21B* KO line #2. Numbers below the images indicate time relative to flower opening. Temporal changes in petal fresh weight (B) and length (C) in wild-type plants, *InMYB21A* KO lines, and *InMYB21B* KO lines. The x-axis indicates time relative to flower opening (0 h). Values are presented as means ± SD (n = 3). Values for wild-type plants are the same as those shown in Figure 1A. (D) Petal diameter in wild-type plants and *InMYB21A* KO line #3. Wild-type plants (n = 54) and *InMYB21A* KO line #3 (n = 90) were compared using Welch’ s *t*-test. (E) Epidermal cell areas on the adaxial surfaces of the limb, ray, and tube in wild-type plants (blue), *InMYB21A* KO line #3 (green), and *InMYB21B*KO line #2 (orange), observed by scanning electron microscopy and quantified. Wild-type plants and each KO line were compared using Welch’ s *t*-test (n = 3; ns, not significant; \**p* < 0. 05, \*\**p* < 0. 01). Temporal changes in glucose (F), sucrose (G), and starch (H) contents in petals of wild-type plants, *InMYB21A* KO lines, and *InMYB21B* KO lines, determined by enzymatic assays. The x-axis indicates time relative to flower opening (0 h). Values are presented as means ± SD in glucose equivalents (nmol Glc/mg FW; n = 3). (I) Cell numbers on the adaxial surfaces of the limb, ray, and tube in wild-type plants (blue), *InMYB21A* KO line #3 (green), and *InMYB21B* KO line #2 (orange). Wild-type plants and each KO line were compared using Welch’ s *t*-test (n = 5; ns, not significant; \**p* < 0. 05, \*\**p* < 0. 01). (J–L) Proportions of cells in the G1, S, and G2/M phases determined by flow cytometric analysis of wild-type plants (J), *InMYB21A* KO line #3 (K), and *InMYB21B* KO line #2 (L). The x-axis indicates time relative to flower opening (0 h), and the y-axis indicates the mean proportion of cells in each cell-cycle phase ± SD (n = 3).

*InMYB21A* KO lines developed and opened their petals similarly to wild-type plants, whereas petal development in *InMYB21B* KO lines began to lag approximately 24 h before flower opening, and their petals ultimately failed to open (Figures 2A and S9). Subsequently, an abscission zone developed at the flower pedicel, resulting in shedding of the entire flower. In *InMYB21A* KO lines, no significant differences from the wild type were observed in changes in petal fresh weight, length, or water content (Figures 2B, 2C, and S10A), or in flower opening and wilting times (Figure S11). In contrast, *InMYB21B* KO lines did not exhibit the rapid increases in petal fresh weight, length, and water content observed in wild-type and *InMYB21A* KO lines from 21 to 3 h before flower opening. Instead, these parameters increased only gradually from 21 h before flower opening to 6 h after flower opening (Figures 2B, 2C, and S10A). These results suggest that *InMYB21B* is required for the rapid petal expansion that begins approximately 21 h before flower opening. In addition, stamen and pistil development was suppressed in *InMYB21B* KO lines from 21 h before flower opening, and the anthers failed to dehisce (Figures S10B, S10C, and S12). Although the distributions of petal diameter largely overlapped, the mean petal diameter of *InMYB21A* KO line #3 was 8.7 mm smaller than that of the wild type (Figure 2D).

### *InMYB21B* promotes petal cell expansion

Organ development is driven by increases in both cell number and cell size. The effects of loss of *InMYB21A* and *InMYB21B* function on petal cell size were therefore investigated. The areas of epidermal cells in the limb, ray, and tube were measured in flowers of wild-type plants, *InMYB21A* KO line #3, and *InMYB21B* KO line #2 at flower opening (Figures 2E and S13A). In *InMYB21A* KO line #3, epidermal cell area in the limb was 0.70-fold (*p* = 0.01) of that in the wild type, whereas no significant differences were observed in the ray or tube. In contrast, in *InMYB21B* KO line #2, epidermal cell areas in the limb, ray, and tube were 0.19-fold (*p* = 2.84E−4), 0.26-fold (*p* = 6.19E−5), and 0.19-fold (*p* = 1.84E−3), respectively, of those in the wild type, and were significantly smaller in all three tissues. These results demonstrate that *InMYB21B* strongly promotes petal cell expansion, whereas loss of *InMYB21A* had a more limited effect on cell size.

In Japanese morning glory petals, an increase in glucose content resulting from starch degradation has been suggested to increase intracellular osmotic pressure and thereby promote petal cell expansion (Figure 1B). To determine whether *InMYB21A* and *InMYB21B* affect sugar metabolism, the contents of glucose, sucrose, and starch in petals of wild-type, *InMYB21A* KO, and *InMYB21B* KO lines were enzymatically quantified at three time points: 39 h before flower opening, 21 h before flower opening, and at flower opening. All values were expressed as glucose equivalents (Figures 2F, 2G, and 2H). In wild-type and *InMYB21A* KO lines, glucose content increased from 39 h before flower opening to flower opening, whereas in *InMYB21B* KO lines, glucose content did not show the marked increase observed in the wild type and *InMYB21A* KO lines (Figure 2F). In contrast, starch content decreased by 87–91% from 21 h before flower opening to flower opening in wild-type and *InMYB21A* KO lines, whereas no apparent decrease was observed in *InMYB21B* KO lines (Figure 2H). These results suggest that *InMYB21B* contributes to petal cell expansion and is required for normal starch degradation and glucose accumulation in petals.

### Loss of *InMYB21B* is associated with increased petal cell number

To evaluate the effects of loss of *InMYB21A* and *InMYB21B* function on petal cell number, cell numbers were determined in flowers of wild-type plants, *InMYB21A* KO line #3, and *InMYB21B* KO line #2 at flower opening (Figures 2I and S13B). In *InMYB21A* KO line #3, no significant differences in cell number were observed relative to the wild type in the limb, ray, or tube. In contrast, *InMYB21B* KO line #2 exhibited significantly increased cell numbers in both the adaxial and abaxial regions of the limb, by 1.47-fold (*p* = 1.31E−4) and 1.50-fold (*p* = 8.42E−5), respectively. Similarly, cell numbers in the adaxial and abaxial regions of the floral tube were increased by 1.64-fold (*p* = 3.43E−4) and 1.87-fold (*p* = 1.81E−4), respectively.

To further evaluate cell-cycle progression in petals, flow cytometric analysis was performed at 12-h intervals from 69 h before flower opening to 3 h after flower opening (Figures 2J–L and S13C–E). In wild-type and *InMYB21A* KO line #3, the proportion of cells in the G2/M phase tended to gradually decrease from 21 h before flower opening (Figures 2J and 2K). In contrast, this decrease was not observed in *InMYB21B* KO line #2, in which the proportion of G2/M-phase cells remained nearly constant from 69 h before flower opening to 3 h after flower opening (Figure 2L). Consequently, the proportion of G2/M-phase cells in petals of *InMYB21B* KO line #2 at 3 h after flower opening was approximately 4.13-fold higher than that in the wild type. These results suggest that the developmental decline or cessation of cell division observed in wild-type petals is impaired in *InMYB21B* KO line #2, resulting in continued cell proliferation and an increase in cell number. Thus, loss of *InMYB21B* is associated with increased petal cell number and prolonged cell-cycle activity during petal development.

### Loss of *InMYB21B* delays transcriptomic progression during petal development

To investigate the effects of *InMYB21B* on the petal transcriptome, the transcriptomes of *InMYB21B* KO lines were compared with those of wild-type and *InMYB21A* KO lines. Transcriptomes were obtained from petals of *InMYB21B* KO line #2 and wild-type plants at 39 h before flower opening, and from petals of *InMYB21A* KO line #3, *InMYB21B* KO line #2, and wild-type plants at 15 h before flower opening. These datasets were combined with the time-series transcriptome dataset obtained from wild-type petals at 3-h intervals from 72 h before flower opening to 12 h after flower opening (Nakagawa et al. 2025), and the resulting datasets were compared by principal component analysis (PCA) (Figure 3A). The transcriptome of *InMYB21B* KO line #2 at 15 h before flower opening was positioned closer in PCA space to that of wild-type petals at 39 h before flower opening than to that of wild-type petals at 15 h before flower opening (Figure 3A). In the wild-type time-series transcriptome, PC1 scores continuously increased from low to high values between 72 h before flower opening and 12 h after flower opening (Figure 3A), indicating that PC1 reflects the developmental progression of petals. These results strongly suggest that the normal transcriptomic progression occurring between 39 and 15 h before flower opening is impaired in *InMYB21B* KO line #2, indicating that loss of *InMYB21B* delays transcriptomic progression during petal development.

**Figure 3.**
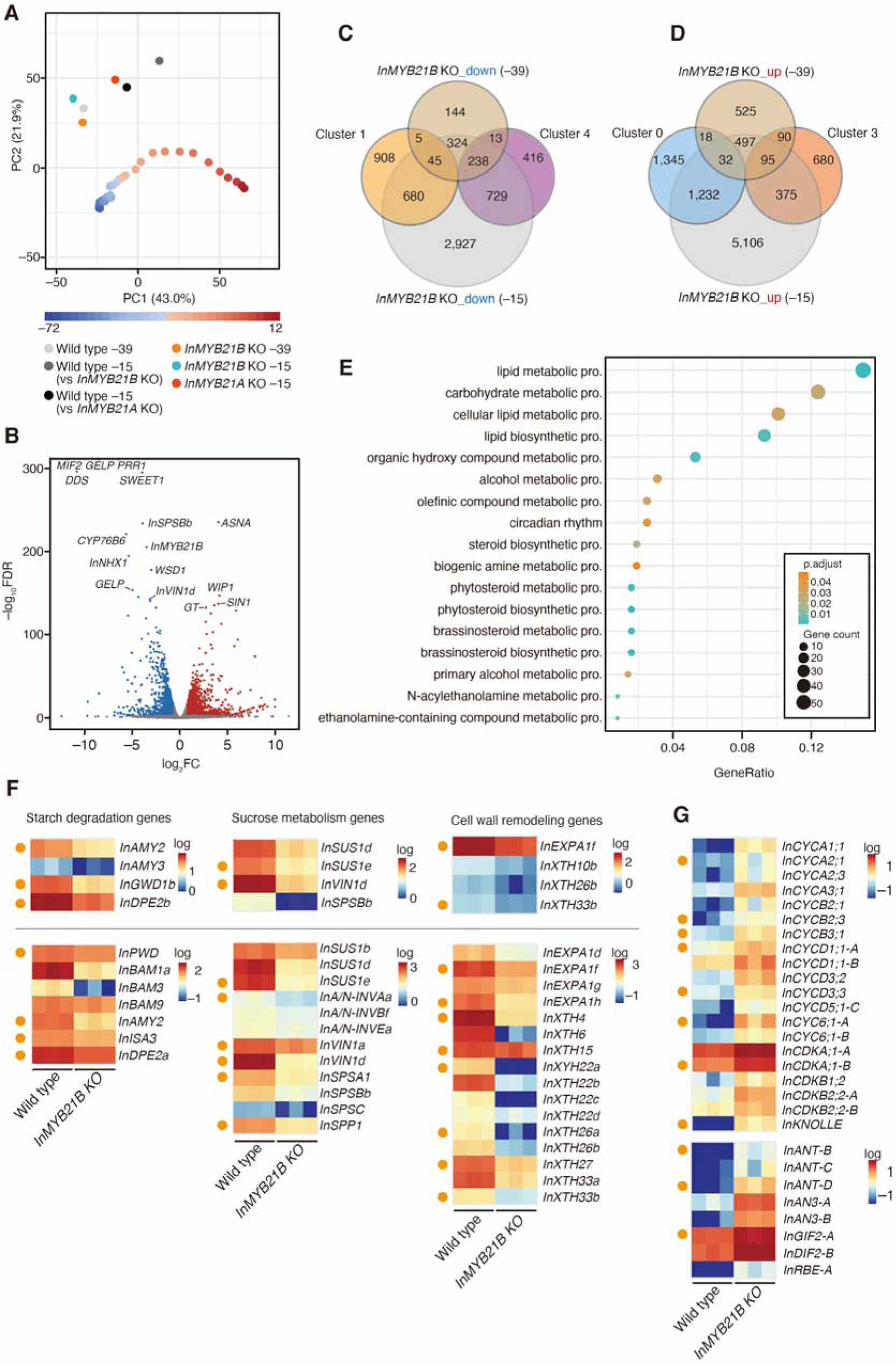
Transcriptome analysis of *InMYB21A* KO and *InMYB21B* KO lines (A) Principal component analysis (PCA) based on transcriptome data. The x- and y-axes indicate principal component 1 and principal component 2, respectively, with the percentages in parentheses indicating the proportions of variance explained. The 29 time-point transcriptome datasets obtained at 3-h intervals from 72 h before flower opening to 12 h after flower opening in our previous study (Nakagawa et al. 2025) are shown as a gradient from blue to red, reflecting developmental progression. (B) Volcano plot showing log2 fold change (log2FC; x-axis) and −log10 false discovery rate (FDR; y-axis) for the transcriptome comparison between *InMYB21B* KO line #2 and wild-type plants at 39 h before flower opening. Downregulated DEGs (log2FC < −1 and −log10FDR > 2) are shown in blue, upregulated DEGs (log2FC > 1 and −log10FDR > 2) in red, and other genes in gray. (C) Venn diagram showing the overlap between genes downregulated in *InMYB21B* KO line #2 at 39 and 15 h before flower opening (Tables S12 and S13) and genes assigned to Clusters 1 and 4 in time-series K-means clustering (Table S8). (D) Venn diagram showing the overlap between genes upregulated in *InMYB21B* KO line #2 at 39 and 15 h before flower opening (Tables S12 and S13) and genes assigned to Clusters 0 and 3 in time-series K-means clustering (Table S8). (E) GO enrichment analysis of DEGs commonly downregulated in *InMYB21B* KO line #2 at 39 and 15 h before flower opening. The significance threshold was set at corrected *p*-value < 0. 05. GeneRatio indicates the proportion of genes in the analyzed DEG set assigned to each GO term. The size of each circle indicates the number of genes, and the color indicates the corrected *p*-value. (F) Heatmap of starch degradation, sucrose metabolism, and cell wall remodeling genes downregulated in *InMYB21B* KO line #2. Expression levels are shown as common logarithms of TPM values, with TPM values of 0 converted to 0. 01 before logarithmic transformation. Each column represents one biological replicate (n = 3). Genes marked with orange circles contain an R2R3-MYB subgroup 19 cis-regulatory sequence within the 1, 000-bp region upstream of the transcription start site. Heatmaps above the black line show DEGs identified at 39 h before flower opening, whereas those below the black line show DEGs identified at 15 h before flower opening. (G) Heatmap of cell cycle-, cell division-, and cell number-promoting genes upregulated in *InMYB21B* KO line #2 at 15 h before flower opening. Expression levels are shown as common logarithms of TPM values, with TPM values of 0 converted to 0. 01 before logarithmic transformation. Each column represents one biological replicate (n = 3). Genes marked with orange circles contain an R2R3-MYB subgroup 19 cis-regulatory sequence within the 1, 000-bp region upstream of the transcription start site.

### Loss of *InMYB21B* affects the expression of more genes than loss of *InMYB21A*

To identify genes whose expression was affected by loss of *InMYB21A* and *InMYB21B*, differentially expressed genes (DEGs) were identified (Tables S11–S13). In petals of *InMYB21A* KO line #3 at 15 h before flower opening, when *InMYB21A* expression began to increase, 100 downregulated and 133 upregulated DEGs were identified relative to the wild type (Figure S14A; Table S11). In contrast, petals of *InMYB21B* KO line #2 were compared with those of the wild type at two time points: at 15 h before flower opening, when *InMYB21B* was highly expressed, and at 39 h before flower opening, during an earlier phase of its expression. At 15 h before flower opening, 4,943 downregulated and 7,337 upregulated DEGs were identified (Figure S14B; Table S13), whereas at 39 h before flower opening, 769 downregulated and 1,257 upregulated DEGs were identified (Figure 3B; Table S12). Among the DEGs detected at 39 h before flower opening, 79% of the downregulated DEGs and 51% of the upregulated DEGs were also detected 24 h later, at 15 h before flower opening (Figure 3C and 3D; Table S14). Furthermore, six downregulated and 33 upregulated DEGs were common to *InMYB21B* KO line #2 at both 39 and 15 h before flower opening and *InMYB21A* KO line #3 at 15 h before flower opening (Figures S14C and S14D).

GO enrichment analysis was performed to investigate the biological functions of the DEGs identified in *InMYB21A* KO line #3. No significantly enriched GO terms were detected among the downregulated DEGs, whereas GO terms associated with organic acid metabolism and lipid metabolism were significantly enriched among the upregulated DEGs (Figure S15A). Based on the predicted functions of the 20 genes included in the enriched GO terms, no apparent association with petal development was identified. Among the six downregulated and 33 upregulated DEGs common to *InMYB21A* KO line #3 at 15 h before flower opening and *InMYB21B* KO line #2 at both 39 and 15 h before flower opening (Figures S14C and S14D), eight were of unknown function, and the predicted functions of the remaining 31 genes did not indicate an apparent association with petal development.

### Loss of *InMYB21B* disrupts gene expression dynamics during petal development

To investigate the temporal expression patterns of DEGs identified in *InMYB21B* KO line #2, the DEGs were compared with the results of time-series K-means clustering (Table S9). Most of the downregulated DEGs in *InMYB21B* KO line #2 belonged to Cluster 4, which exhibited peak expression around flower opening, or Cluster 1, which exhibited peak expression after flower opening. Cluster 4 contained 251 DEGs identified at 39 h before flower opening and 967 DEGs identified at 15 h before flower opening, accounting for 18% and 69%, respectively, of all genes assigned to Cluster 4 (Figure 3C; Table S9). Cluster 1 contained 725 DEGs identified at 15 h before flower opening, accounting for 44% of all genes assigned to Cluster 1 (Figure 3C; Table S9). In contrast, the 185 DEGs upregulated at 39 h before flower opening were most strongly represented in Cluster 3, which exhibited a gradual decrease in expression and accounted for 14.9% of all genes assigned to Cluster 3 (Figure 3D; Table S9). Among the DEGs upregulated at 15 h before flower opening, 1,264 belonged to Cluster 0, which also exhibited a gradual decrease in expression and accounted for 48% of all genes assigned to Cluster 0 (Figure 3D; Table S9). These results suggest that loss of *InMYB21B* is associated with reduced expression of genes that exhibit transient (Cluster 4) or sustained (Cluster 1) increases in expression during petal development from 72 h before flower opening to 12 h after flower opening, and with increased expression of genes that exhibit decreasing expression patterns (Clusters 0 and 3).

### Loss of *InMYB21B* alters expression of genes associated with petal growth and sugar content

The relationship between DEGs identified in *InMYB21B* KO line #2 and the co-expression modules associated with petal phenotypes was examined. Genes belonging to the cyan, darkred, green, and grey60 co-expression modules, which were positively correlated with petal fresh weight, petal length, glucose content, and sucrose content, were frequently identified among the downregulated DEGs at 15 h before flower opening, accounting for 76%, 70%, 30%, and 42% of the genes in these modules, respectively (Figure 1F). In contrast, genes belonging to the darkgreen, lightcyan, turquoise, darkturquoise, and lightyellow modules, which were positively correlated with starch content, were frequently identified among the upregulated DEGs at 15 h before flower opening, accounting for 67%, 42%, 38%, 39%, and 42%, respectively, of the genes in these modules (Figure 1F). These results suggest that loss of *InMYB21B* was associated with reduced expression of genes positively correlated with petal fresh weight, petal length, glucose content, and sucrose content, and increased expression of genes positively correlated with starch content.

### Loss of *InMYB21B* is associated with reduced expression of genes involved in sugar metabolism, cell wall remodeling, and water transport

To investigate the functions of genes whose expression was affected by loss of *InMYB21B*, GO enrichment analysis was performed for DEGs identified in *InMYB21B* KO line #2. Among the downregulated DEGs, 20 GO terms were significantly enriched at 39 h before flower opening (Figure S15B), and 108 GO terms were significantly enriched at 15 h before flower opening (Table S16). Furthermore, 17 GO terms were commonly enriched among the DEGs downregulated at both 39 and 15 h before flower opening, including terms significantly associated with lipid metabolism and sugar metabolism (Figure 3E).

Among the sugar metabolism-related genes commonly downregulated at both 39 and 15 h before flower opening were the starch degradation gene *InBAM1a* and the sucrose metabolism genes *InVIN1a*, *InVIN1d*, *InSUS1b*, and *InSUS1d*, which may contribute to the increase in osmotic pressure during flower opening (Figure 3F). GO terms related to cell wall remodeling and cell wall metabolism were also enriched among the DEGs downregulated at 15 h before flower opening. These DEGs included *InXTH4*, *InXTH6*, and *InEXPA1f*, which may contribute to cell wall remodeling during flower opening (Figure 3F).

In addition, 19 genes encoding aquaporins were downregulated in *InMYB21B* KO line #2 at 15 h before flower opening (Figure S16). Among these, *InPIP1;2*, which encodes a plasma membrane aquaporin, reached its expression peak at flower opening in Japanese morning glory and exhibited an exceptionally high FPKM value of approximately 15.5K, suggesting that it may contribute to water transport in petals (Nakagawa et al. 2025). Taken together, these results suggest that *InMYB21B* promotes petal cell expansion, possibly through changes in sugar metabolism and the expression of genes involved in cell wall remodeling and water transport.

### Loss of *InMYB21B* is associated with increased expression of genes involved in the cell cycle and cell division

GO enrichment analysis was performed for the DEGs upregulated in *InMYB21B* KO line #2. Unlike the downregulated DEGs, the upregulated DEGs showed no significant enrichment of GO terms related to sugar metabolism or cell wall processes (Figure S15; Tables S17 and S18). A total of 102 GO terms were significantly enriched at 39 h before flower opening (Table S17), and 149 GO terms were significantly enriched at 15 h before flower opening (Table S18). Among the DEGs commonly upregulated at 39 and 15 h before flower opening, 20 GO terms associated with stress responses, lipid metabolism, organic acid metabolism, and photosynthesis were significantly enriched (Figure S15C).

Among the GO terms enriched at 15 h before flower opening were “cell cycle” and “cell division.” Examination of the genes associated with these GO terms identified 19 cell cycle-related genes and *InKNOLLE*, a gene known as a marker of cell division (Figure 3G). The expression levels of 43 cell cycle-related genes and *InKNOLLE* were therefore compared among wild-type plants at 39 h before flower opening and *InMYB21B* KO line #2 at 39 and 15 h before flower opening using a heatmap (Figure S17). No marked differences in expression levels were observed among these samples. These results suggest that the expression of cell cycle- and cell division-related genes remained at levels characteristic of an earlier developmental stage in *InMYB21B* KO petals.

In Arabidopsis, several genes have been reported to regulate petal cell number by controlling the activity and duration of cell division (Krizek and Anderson 2013). To investigate whether loss of *InMYB21B* affects the expression of these genes, their Japanese morning glory orthologs were identified (Table S19), and their expression patterns were examined by DEG analysis (Figure 3G). *AINTEGUMENTA* (*ANT*), which encodes an AP2/ERF-type transcription factor, promotes petal cell proliferation in response to auxin signaling (Krizek 1999; Krizek and Anderson 2013). Three Japanese morning glory orthologs, *InANT-B*, *InANT-C*, and *InANT-D*, were upregulated in *InMYB21B* KO line #2 (Figure 3G). *ANGUSTIFOLIA3* (*AN3*), which functions as a cell proliferation factor (Lee et al. 2009; Zhang et al. 2019), also has two Japanese morning glory orthologs, *InAN3-A* and *InAN3-B*, both of which were upregulated. *RABBIT EARS* (*RBE*), which encodes a C2H2 zinc-finger transcriptional repressor, promotes cell division activity and prolongs the period during which cell division remains active in Arabidopsis petals, thereby contributing to an increase in petal cell number (Huang and Irish 2015). The Japanese morning glory ortholog *InRBE-A* was also upregulated. These results suggest that loss of *InMYB21B* was associated with continued cell proliferation and increased expression of genes involved in promoting cell proliferation.

### Loss of *InMYB21B* is associated with increased expression of jasmonate biosynthetic genes

R2R3-MYB SG19 transcription factors negatively regulate jasmonate biosynthesis in Arabidopsis but positively regulate it in tomato and rice (Gou et al. 2024; Huang et al. 2017; Reeves et al. 2012; Schubert et al. 2019). The involvement of *InMYB21B* in jasmonate biosynthesis in Japanese morning glory petals was therefore investigated. Orthologs of genes involved in jasmonate biosynthesis and signaling in Japanese morning glory were identified (Tables S20 and S21), and their expression patterns in petals were visualized using a heatmap (Figure S18). Among these genes, *InLOX5A*, *InLOX5B*, and *InAOC1A* exhibited high expression levels, which decreased after 3 h before flower opening. All three genes were upregulated in *InMYB21B* KO line #2 at 39 h before flower opening (Table S20). These results suggest that, as in Arabidopsis, loss of *InMYB21B* was associated with increased expression of jasmonate biosynthetic genes in Japanese morning glory.

## DISCUSSION

In Japanese morning glory, petal development is driven by both cell proliferation and cell expansion until approximately 48 h before flower opening, whereas development from this point until flower opening is primarily driven by cell expansion (Nakagawa et al. 2025). Notably, *InMYB21B* expression begins to increase at approximately the same developmental stage. In the present study, loss of *InMYB21B* markedly impaired petal cell expansion and prevented flower opening. This phenotype was accompanied by impaired starch degradation and glucose accumulation and by delayed transcriptomic progression during late petal development. Together, these findings identify InMYB21B as an important regulator of the rapid petal expansion that precedes flower opening.

### Function of *InMYB21B* in petals

Petal cell expansion is thought to involve an increase in osmotic pressure resulting from the degradation of polysaccharides and cell wall loosening (van Doorn and Kamdee 2014; van Doorn and Van Meeteren 2003). In Japanese morning glory, petals begin to expand rapidly approximately 21 h before flower opening (Figure 1A). This rapid expansion was preceded by increases in glucose and sucrose contents from approximately 27 h before flower opening, followed by a decrease in starch content from 21 h before flower opening (Figure 1B). These changes are consistent with the possibility that increased osmotic pressure associated with sugar accumulation contributes to petal cell expansion in Japanese morning glory. During this period, the expression levels of *InBAM1a*, involved in starch degradation, *InVIN1d* and *InSUS1d*, involved in sucrose metabolism, and *InXTH4*, *InXTH6*, and *InEXPA1f*, involved in cell wall remodeling, increased. These genes may therefore contribute to petal cell expansion during flower opening.

To investigate the relationships between the petal transcriptome from 72 h before flower opening to 12 h after flower opening and petal fresh weight, petal length, and glucose, sucrose, and starch contents, weighted gene co-expression network analysis (WGCNA) was performed (Figure 1E). This analysis identified *InMYB21B*, which was specifically expressed in floral organs, as a hub gene in the cyan co-expression module, which showed the strongest association with glucose content. Furthermore, genome-edited *InMYB21B* KO lines exhibited suppressed rapid petal expansion beginning approximately 21 h before flower opening, resulting in failure of the petals to open (Figures 2A–2C; Figure S9). Flow cytometric analysis suggested that cell division remained active in *InMYB21B* KO petals even after flower opening. In addition, DEG analysis of petals at 15 h before flower opening revealed increased expression of genes associated with the cell cycle and cell division. These results suggest that loss of *InMYB21B* is associated with prolonged cell-cycle activity during petal development.

Cell size analysis further demonstrated that cell expansion was impaired in *InMYB21B* KO lines. DEG analysis revealed reduced expression of *InBAM1a*, which is involved in starch degradation; *InVIN1d* and *InSUS1d*, which are involved in sucrose metabolism; *InXTH4*, *InXTH6*, and *InEXPA1f*, which are involved in cell wall remodeling; and *InPIP1;2*, which is associated with water transport (Figures 3F and S16). Comparison of the DEGs with the co-expression modules identified by WGCNA further suggested that loss of *InMYB21B* was associated with reduced expression of genes positively correlated with petal fresh weight, petal length, and glucose and sucrose contents, and increased expression of genes positively correlated with starch content. These findings indicate that *InMYB21B* promotes petal cell expansion, possibly through changes in sugar metabolism and the expression of genes involved in water transport and cell wall remodeling. They also support the importance of *InMYB21B* during the later phase of petal development, when cell expansion predominates. Consistent with this interpretation, the transcriptomic profiles of *InMYB21B* KO petals collected 15 h before flower opening, as assessed by PCA, were more similar to those of wild-type petals at 39 h before flower opening than to those at 15 h before flower opening, suggesting that the developmental progression of the KO lines may be delayed.

In addition to the genes described above, the expression levels of numerous other genes were altered in *InMYB21B* KO petals. At 15 h before flower opening, 4,943 downregulated DEGs and 7,337 upregulated DEGs were identified. Comparison of these DEGs with the clusters identified by time-series K-means clustering based on their temporal expression patterns further suggested that loss of *InMYB21B* was associated with reduced expression of genes showing increasing expression patterns (Clusters 1 and 4) and increased expression of genes showing decreasing expression patterns (Clusters 0 and 3) during petal development from 72 h before flower opening to 12 h after flower opening. GO enrichment analysis further revealed significant enrichment of GO terms associated with lipid metabolism and other biological processes. These results suggest that loss of I*nMYB21B* causes broad transcriptomic changes during petal development, affecting not only genes involved in sugar metabolism, water transport, and cell wall remodeling, but also those involved in processes such as lipid metabolism.

R2R3-MYB SG19 transcription factors have been reported to recognize cis-regulatory sequences related to [G/A]TT[A/T]GG[T/C], suggesting that InMYB21B may recognize a similar sequence. However, this sequence occurs relatively frequently by chance, with an expected frequency of approximately once every 2,048 bp. Assuming a promoter length of 1,000 bp, there is approximately a 38.6% probability that this sequence will occur within a promoter by chance. Although this cis-regulatory sequence is present in the promoter regions of some genes identified as DEGs in *InMYB21B* KO lines, it is difficult to statistically determine whether these occurrences represent functional binding sites or chance occurrences. Therefore, further experiments, such as chromatin immunoprecipitation followed by sequencing (ChIP-seq) or DNA-binding assays, will be required to determine the direct targets of *InMYB21B*.

### Relationship between *InMYB21B*, its orthologs, and phytohormones

R2R3-MYB SG19 transcription factors have diverse roles in flower development and maturation. In *P. axillaris*, partial loss of EOB2 function results in reduced petal cell size and impaired starch degradation, indicating that EOB2 is involved in petal maturation and primary metabolism (Chopy et al. 2023). In rice, OsMYB8 promotes floret opening by directly activating *OsJAR1*, thereby affecting JA-Ile accumulation and the expression of genes associated with lodicule hydration and cell wall remodeling (Gou et al. 2024). The present study extends these findings by showing that loss of *InMYB21B* causes a pronounced defect in petal cell expansion together with impaired starch degradation and glucose accumulation and delayed transcriptomic progression during petal development. In addition, loss of *InMYB21B* was associated with increased petal cell number and prolonged cell-cycle activity, providing evidence that its effects on petal development extend beyond cell expansion.

The cessation of cell division during plant organ development is thought to be regulated by a network involving multiple phytohormones, including gibberellins, cytokinins, and brassinosteroids (Achard et al. 2009; Jing et al. 2023; Nelissen et al. 2012; Skalak et al. 2019; Zhiponova et al. 2013). Recent studies in rose have further shown that cytokinin-responsive transcriptional modules regulate the duration and activity of petal cell proliferation (Jin et al. 2025; Wang et al. 2024). In Arabidopsis, *ANT* promotes cell proliferation (Krizek 1999; Krizek and Anderson 2013). In the present study, loss of *InMYB21B* was associated with increased expression of Arabidopsis ANT orthologs in Japanese morning glory. This association suggests a possible link between *InMYB21B* and ANT-mediated regulation of cell proliferation, although the underlying regulatory relationship remains to be determined.

In addition, loss of *InMYB21B* was associated with increased expression of genes involved in jasmonate biosynthesis, including *InLOX5A*, *InLOX5B*, and *InAOC1A*. R2R3-MYB SG19 transcription factors are also known to regulate jasmonate biosynthesis in Arabidopsis, tomato, and rice. Although tomato and Japanese morning glory both belong to the order Solanales, *SlMYB21* in tomato positively regulates jasmonate biosynthesis (Schubert et al. 2019), indicating that R2R3-MYB SG19 transcription factors can exert opposite effects on jasmonate biosynthesis even among members of the same plant order. Jasmonate is also known to suppress cell proliferation in Arabidopsis leaves by regulating the expression of cell cycle-related genes (Noir et al. 2013; Zhang and Turner 2008). However, the effects of jasmonate on cell proliferation may depend on developmental and species contexts. In rose petals, RhMYC2 contributes to maintaining the duration of the cell division phase through regulation of cytokinin homeostasis (Gong et al. 2024). Because jasmonate levels were not measured in the present study, the continued cell proliferation observed in *InMYB21B* KO lines cannot be attributed to changes in jasmonate levels. Moreover, the increased expression of jasmonate biosynthetic genes in *InMYB21B* KO lines does not support a simple model in which loss of *InMYB21B* reduces jasmonate levels and thereby promotes continued cell proliferation. Further analyses of jasmonate levels and jasmonate signaling will be necessary to determine whether the altered expression of jasmonate biosynthetic genes contributes to the regulation of cell proliferation by *InMYB21B*. Nevertheless, how R2R3-MYB SG19 transcription factors, including *InMYB21B*, interact with phytohormone networks to regulate floral organ development remains largely unknown and warrants further investigation.

### Function of *InMYB21B* in floral organs other than petals

R2R3-MYB SG19 transcription factors are involved not only in petal development but also in pistil and stamen development (Chopy et al. 2023; Colquhoun et al. 2011; Mandaokar et al. 2006; Niwa et al. 2018; Reeves et al. 2012; Yarahmadov et al. 2020). In Arabidopsis, *MYB21* and *MYB24* are involved in pistil elongation and anther dehiscence, and *myb21* loss-of-function mutants exhibit male sterility. However, because seeds can be produced when wild-type pollen is applied to the stigmas of *myb21* mutants, female fertility is largely retained in Arabidopsis *myb21* and *myb21 myb24* mutants (Mandaokar et al. 2006; Reeves et al. 2012). In tomato, *Slmyb21* loss-of-function mutants exhibit female sterility associated with defective ovule development, whereas stamen development and pollen viability are largely unaffected (Schubert et al. 2019). In *P. axillaris*, an *EOB2* mutant lacking the transcriptional activation domain shows defective stigma maturation and reduced seed set, but no apparent stamen or pollen phenotype (Chopy et al. 2023). In rice, *Osmyb8* loss-of-function plants have been reported to exhibit neither male nor female sterility (Gou et al. 2024).

In Japanese morning glory, *InMYB21B* was expressed not only in petals but also in stigmas and anthers. In *InMYB21B* KO lines, stamen and pistil development was suppressed from approximately 21 h before flower opening, and anther dehiscence did not occur (Figures S10B, S10C, and S12). Reciprocal crossing experiments with wild-type plants further demonstrated that the KO lines were both female- and male-sterile. In contrast, no apparent fertility defects were observed in *InMYB21A* KO lines. Although Japanese morning glory, tomato, and petunia all belong to the order Solanales, these findings indicate substantial interspecific diversity in the roles of R2R3-MYB SG19 transcription factors in male and female fertility. Elucidating the molecular basis of these interspecific differences will be important for understanding the evolutionary diversification of floral organ development. Comparative studies of R2R3-MYB SG19 transcription factors across diverse plant species will therefore be an important direction for future research.

### Function of *InMYB21A*

*InMYB21A* KO lines showed no obvious abnormalities in petal development and underwent normal flower opening. Furthermore, no significant differences from the wild type were observed in petal fresh weight, petal length, or water content. Although *InMYB21A* KO line #3 showed a statistically significant reduction in petal diameter, the distributions largely overlapped and no significant differences were detected in petal fresh weight, length, water content, or flower opening time (Figures 2B–2D, S10A, and S11). These results suggest that *InMYB21A* may have a limited role in promoting cell expansion during petal development.

*InMYB21A* contains a single-amino-acid insertion within a transcriptional activation domain that is conserved across plant species (Figure S5A), suggesting that its transcriptional activation activity may be reduced or lost. However, this amino acid insertion is conserved within the genus Ipomoea, suggesting that *InMYB21A* may retain a functional role. Furthermore, 233 DEGs were identified in petals of *InMYB21A* KO line #3 at 15 h before flower opening, indicating that *InMYB21A* contributes to transcriptional regulation. However, the predicted functions of these DEGs did not provide sufficient information to identify the biological processes regulated by *InMYB21A*. Further analyses using additional independent KO lines will be necessary to clarify the functions of *InMYB21A* in petal development.

### Future perspectives

The present study demonstrates that I*nMYB21B* is required for normal flower opening but does not establish whether InMYB21B regulates the timing of flower opening. In rice, differences in flower opening time between indica and japonica varieties are attributable to differences in the expression of the R2R3-MYB SG19 transcription factor gene *OsMYB8*, associated with promoter polymorphisms (Gou et al. 2024). These findings raise the possibility that *InMYB21B* expression may also contribute to variation in flower opening time in Japanese morning glory.

Japanese morning glory accessions exhibit considerable variation in flower opening time. Comparative analyses of *InMYB21B* promoter sequences and expression patterns among accessions with different flower opening times may help clarify the relationship between *InMYB21B* and the regulation of flower opening time. In addition, flower opening in Japanese morning glory is controlled by the circadian clock (Kaihara and Takimoto 1979), and clock genes exhibiting circadian expression patterns in petals were identified by Nakagawa et al. (2025). Investigating whether the expression of *InMYB21B* is regulated by these clock genes may clarify whether *InMYB21B* serves as a molecular link between the circadian clock and flower opening.

## MATERIALS AND METHODS

### Plant materials

Plants used in this study were obtained by self-pollination of a next-generation plant derived from a single individual of the Tokyo Kokei Standard line used for whole-genome sequencing (Hoshino et al. 2016). Plants were grown in commercially available potting soil in 5-inch pots in an indoor growth room under a 13-h light/11-h dark cycle at 25°C. For sampling during the dark period, a green light was used to prevent exposure of the plants to other wavelengths of light. The flower opening time was defined as 08:00 on the day of anthesis. After petal sampling, the fresh weight of the petals was measured under illuminated conditions. The petals, stamens, and pistils were photographed using a digital camera (Nikon Z50; Nikon, Japan), and their lengths were measured from the digital images using ImageJ Fiji (Schindelin et al. 2012). Petal diameter was measured in the same manner. To determine petal dry weight, the petals were dried at 50°C for 24 h in a constant-temperature drying oven (MOV-212F; Sanyo Electric, Japan). Water content was calculated by subtracting dry weight from fresh weight.

### Generation of genome-edited plants

Binary vectors expressing both CRISPR/Cas9 and sgRNAs were constructed based on pDE_CAS9_KAN (Fauser et al. 2014). Target sequences for sgRNAs specific to the target genes were designed using the CRISPRdirect web-based tool (http://crispr.dbcls.jp/), with two target sites selected for each of *InMYB21A* and *InMYB21B* (Figures S6 and S7). The target sites were located in exons 1 and 2 of *InMYB21A* and were both located in exon 2 of *InMYB21B*. Target sequences for the sgRNAs were introduced into pMR217 and pMR218 by PCR amplification using these vectors as templates and target-specific primers together with the common primer pMR-Fw1 (Table S22), followed by circularization of the PCR products. The resulting sgRNA expression cassettes were subsequently cloned into the pDE_CAS9_KAN destination vector by an LR reaction using Gateway LR Clonase II (Thermo Fisher Scientific, USA). The resulting binary vectors were introduced into *Agrobacterium tumefaciens* strain EHA105. The Tokyo Kokei Standard line was transformed with the binary vectors according to the method described by Takatori et al. (2015). Genomic DNA was extracted from regenerated plants using PI-480 (Kurabo, Japan). DNA fragments containing *InMYB21A* and *InMYB21B* were amplified by PCR and analyzed using MultiNA (Shimadzu Corporation, Japan), followed by DNA sequencing. Plants carrying genome-edited mutations in either *InMYB21A* or *InMYB21B* were selected for further analyses.

### Flower opening and petal development

To observe the flower opening process, time-lapse images were captured at 30-min intervals from 24 h before the defined flower opening time until flower opening using a digital camera (Nikon D5100; Nikon, Japan). To quantitatively evaluate the flower opening process, the corolla area was also measured. Time-lapse images were captured at 30-min intervals from 15 h before the defined flower opening time (17:00 on the day before flower opening) to 12 h after flower opening (20:00 on the day of flower opening) using a digital camera (Nikon D5100; Nikon, Japan). The camera was positioned perpendicular to the petals. For imaging under dark conditions, the built-in flash of the camera was used. Following the method described by Shinozaki et al. (2011), only the colored region of the petals (corolla) was extracted from the images, and the relative corolla area at each time point was calculated by dividing the corolla area by the maximum corolla area of each flower.

The cell cycle was analyzed using a flow cytometer. Approximately 5-mm-square sections of the limb, tube, and ray were excised, and nuclei were isolated and stained with DAPI according to the manufacturer’s instructions for the Quantum Stain NA UV 2 Kit (Partec, Germany). The stained nuclei were analyzed using a CyFlow SL flow cytometer and Partec FloMax software (both from Partec, Germany), and the proportions of nuclei in each cell-cycle phase were calculated.

Petal cell area was measured from images obtained using a tabletop scanning electron microscope (TM4000Plus; Hitachi High-Tech, Japan) and analyzed using ImageJ Fiji (Schindelin et al. 2012). For the limb, tube, and ray, the areas of 100 cells from three flowers were measured.

The numbers of cells in the limb, ray, and tube were estimated by calculating the cell density per unit area and multiplying it by the surface area of each tissue. Mean cell density per unit area was calculated from three image datasets. To determine the surface area of each tissue, the tissue was spread by attaching it to adhesive tape and photographed using a digital camera (Nikon Z50; Nikon, Japan). Surface area was then calculated from the digital images using ImageJ Fiji (Schindelin et al. 2012).

### RNA extraction and sequencing

Petals were sampled from *InMYB21A* KO line #3 at 15 h before flower opening and from *InMYB21B* KO line #2 at 39 and 15 h before flower opening. Petals from wild-type plants were also sampled at the corresponding time points. Three biological replicates were collected for each genotype and time point. Total RNA was extracted from the petals using the Maxwell RSC RNA Plant System (Promega, USA), with the following modifications to the manufacturer’s protocol. Briefly, 70–200 mg of frozen, pulverized petal tissue was transferred to 500 µL of homogenization buffer supplemented with 10 µL of thioglycerol and further homogenized with a mortar and pestle at room temperature. The entire homogenate was then centrifuged, and the supernatant was added to the lysis buffer. The extracted RNA was quantified using the Qubit RNA BR Assay Kit and a Qubit fluorometer (both from Thermo Fisher Scientific, USA).

RNA extracted from *InMYB21A* KO line #3 at 15 h before flower opening, *InMYB21B* KO line #2 at 39 h before flower opening, and the corresponding wild-type petals was subjected to next-generation sequencing (NGS) analysis by Genome Read Inc. (Japan). mRNA was purified using the KAPA mRNA Capture Kit (Illumina, USA), and sequencing libraries were prepared using the MGIEasy RNA Directional Library Prep Set (MGI, China). Sequencing was performed on a DNBSEQ-G400RS platform (MGI, China) using 150-bp paired-end reads. RNA extracted from the petals of *InMYB21B* KO line #2 and wild-type plants at 15 h before flower opening was subjected to library preparation and sequencing by Novogene (China). mRNA was purified using poly-T-conjugated magnetic beads, and sequencing libraries were prepared using the NEBNext Ultra II Directional RNA Library Prep Kit (New England Biolabs, USA). Sequencing was performed on a DNBSEQ-T7 platform (BGI, China) using 150-bp paired-end reads.

The obtained sequence data were trimmed using fastp version 0.23.2 (Chen et al. 2018), and the trimmed reads were mapped to the mRNA sequences reported by Hoshino et al. (2016) using kallisto version 0.46.2 (Bray et al. 2016). Gene expression levels were quantified as transcripts per million (TPM), and estimated counts generated by kallisto were used as input for differential expression analysis with edgeR. Reads were also mapped to transcript sequences including splice variants, and the estimated counts of individual splice variants were summed for each gene before edgeR analysis. Principal component analysis (PCA) was performed using the prcomp function in R after log10 transformation of TPM values after adding 1 to each value. Plots were generated using ggplot2 version 3.4.2.

Differentially expressed genes (DEGs) were identified using the R package edgeR version 3.36.0 (Robinson et al. 2010) based on the estimated counts described above. Genes with FDR < 0.01 and |log2FC| > 1 were considered differentially expressed. Volcano plots were generated using the tidyverse version 2.0.0 and ggrepel version 0.9.3 R packages based on the FDR and logFC values calculated by edgeR. Heatmaps showing gene expression levels were generated from log10-transformed TPM values using ggplot2 version 3.4.2 and RColorBrewer version 1.1-3. TPM values of zero were replaced with 0.01 before log10 transformation.

Gene Ontology (GO) enrichment analysis was performed using BiNGO: Biological Network Gene Ontology version 3.0.3 (Maere et al. 2005), with a significance threshold of corrected *p*-value < 0.05. GO terms were obtained from Hoshino et al. (2016), and the ontology file was obtained from the Gene Ontology Consortium (https://geneontology.org/docs/download-ontology/). Statistical results calculated by BiNGO were visualized using ggplot2 version 3.4.2.

### Bioinformatic analysis

The orthologous gene dataset generated using OrthoFinder and reported by Nakagawa et al. (2025) was used for the analysis.

Weighted gene co-expression network analysis (WGCNA) was performed using the R package WGCNA version 1.72-1 (Langfelder and Horvath 2008). The gene expression dataset reported by Nakagawa et al. (2025) was used for the analysis. Genes with a mean FPKM value of < 10 were excluded before analysis, and expression modules were constructed using the blockwiseModules function. The main parameters were set as follows: power = 10, TOMType = “unsigned”, mergeCutHeight = 0.25, and minModuleSize = 30. Pearson correlation analysis was used to calculate correlation coefficients between each co-expression module and petal weight, petal length, glucose content, sucrose content, and starch content. Networks among the modules were visualized using Cytoscape version 3.10.0 (Shannon et al. 2003).

Time-series K-means clustering analysis was performed using the Python library tslearn based on the dataset reported by Nakagawa et al. (2025). The 10,000 genes with the highest mean FPKM values were used for clustering. The elbow method was used to determine the appropriate number of clusters, and the number of clusters was set to six. FPKM values for each gene were standardized to z-scores and clustered using the TimeSeriesKMeans function. Clustering results were visualized using the R package ggplot2 version 3.4.2. These criteria are summarized in Table S15.

For Harr plots, 1,000-bp genomic sequences upstream of the transcription start sites of *InMYB21A*and *InMYB21B* were extracted based on the genome sequence reported by Hoshino et al. (2016) and analyzed using GENETYX-MAC version 21.0.1.

To identify putative MYB-binding motifs in the promoter regions of the DEGs, 1,000-bp upstream sequences from the transcription start sites (TSSs) were extracted from the reference genome and searched for the consensus sequence [G/A]TT[A/T]GG[T/C]. Both the forward strand and its reverse complement were searched for the motif.

### Sugar and starch analysis

Glucose, sucrose, and starch contents were determined with modifications to previously described methods (Araya et al. 2006; Sugiura et al. 2015). Collected petal samples were frozen in liquid nitrogen and ground using a TissueLyser II (QIAGEN, Germany). Soluble sugars were extracted by adding 0.5 mL of 80% ethanol to the powdered samples and incubating them at 80°C for 30 min. This extraction procedure was repeated twice. The resulting pellet was used for starch determination. The supernatants obtained after extraction were concentrated using a Savant SPD1010 SpeedVac (Thermo Fisher Scientific, USA) at 55°C for 2.5 h. After concentration, 150 µL each of distilled water and chloroform were added, and the aqueous phase was collected to separate soluble sugars from soluble proteins. The aqueous phase was used for glucose and sucrose determination.

For sucrose determination, 20 µL of the supernatant was mixed with 20 µL of yeast-derived invertase solution (FUJIFILM Wako Pure Chemical Corporation, Japan) and incubated at 25°C for 60 min to hydrolyze sucrose into glucose and fructose. For starch determination, 0.5 mL of distilled water was added to the pellet, followed by incubation at 100°C for 60 min. After cooling the samples to 55°C, 0.5 mL of amyloglucosidase solution (35 units/mL), prepared by dissolving amyloglucosidase (Sigma-Aldrich, USA) in 100 mM sodium acetate buffer (pH 4.5), was added. The samples were then incubated at 55°C for 60 min to hydrolyze starch into glucose.

Glucose content was quantified using the Glucose CII-Test Wako kit (FUJIFILM Wako Pure Chemical Corporation, Japan) and a microplate reader (SH-9000Lab; Hitachi, Japan). Sucrose content was calculated by subtracting the glucose content measured in the glucose assay sample from the glucose content measured in the sucrose assay sample. Starch content was calculated based on the amount of glucose generated by enzymatic hydrolysis of starch.

### Phylogenetic analysis

Multiple sequence alignment was performed using the high-accuracy E-INS-i option of MAFFT version 7.505 (Katoh and Standley 2013), implemented in Jalview version 2.10.5 (Waterhouse et al. 2009). Gapped regions were removed using trimAl version 1.5.rev0 (Capella-Gutierrez et al. 2009). A phylogenetic tree was constructed using the maximum-likelihood method implemented in MEGA11 (Tamura et al. 2021), and node reliability was assessed using 1,000 bootstrap replicates. The amino acid substitution model was based on the Jones–Taylor–Thornton (JTT) matrix (Jones et al. 1992), incorporating a gamma distribution among sites. The subtree pruning and regrafting (SPR) method, with a search level of 5, was used as the heuristic method. The resulting phylogenetic tree was visualized using Interactive Tree Of Life (iTOL) version 6 (Letunic and Bork 2024).

### Statistical analysis

Normality was assessed using the Anderson–Darling test, with *p* > 0.05 considered indicative of a normal distribution. For comparisons between two groups, when the data were considered normally distributed, Welch’s *t*-test was performed using the t.test function implemented in R, with *p* < 0.05 considered statistically significant. When the data did not meet the normality criterion, the Brunner–Munzel test was performed using the R package lawstat version 3.6 (Hui et al. 2008), with *p* < 0.05 considered statistically significant.

## ACKNOWLEDGEMENTS

The authors thank Sachiko Tanaka for her early work on *InMYB21A*, which contributed to the initiation of this study, and Shizuka Koshimizu for her guidance on bioinformatics analysis. The authors also thank Seiichi Toki and Masaki Endo for providing the CRISPR/Cas9 vectors, Masayoshi Kawaguchi and Hiroshi Ito for valuable discussions, Kensuke Kawade for support with sugar analysis, and Kazuyo Ito, Tomoyo Takeuchi, Saki Kawada, Kiyoko Kuzunishi, and Naoko Koyama for technical support. This study utilized equipment at the Model Organisms Facility, Data Integration and Analysis Facility, Trans-Omics Facility, and Optics and Imaging Facility, NIBB Trans-Scale Biology Center.

