## Supplemental Figures S1-S18 for "*InMYB21B* Promotes Petal Cell Expansion and Flower Opening in Japanese Morning Glory (*Ipomoea nil*)"

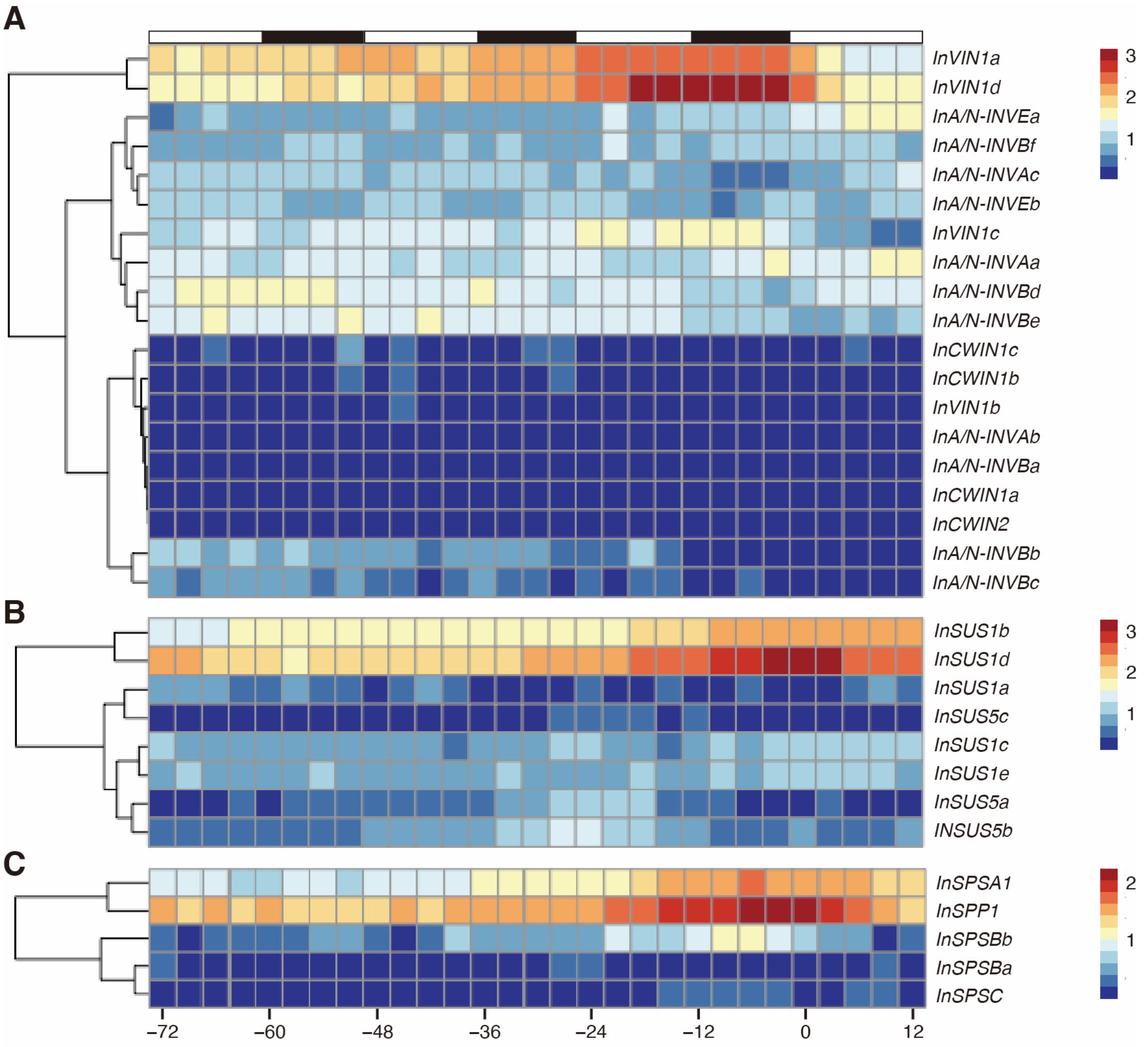


**Figure S1.** Heatmap analysis of the temporal expression patterns of genes involved in sugar metabolism

Heatmaps showing the expression patterns of *invertase* genes (**A**), *sucrose synthase* genes (**B**), and genes involved in sucrose biosynthesis (**C**). The gene lists are provided in Tables S2, S3, and S4, respectively. Expression levels are shown as log10(FPKM + 1). The x-axis indicates time relative to flower opening (0 h). The bars at the top indicate light conditions, with white and black representing light and dark periods, respectively.

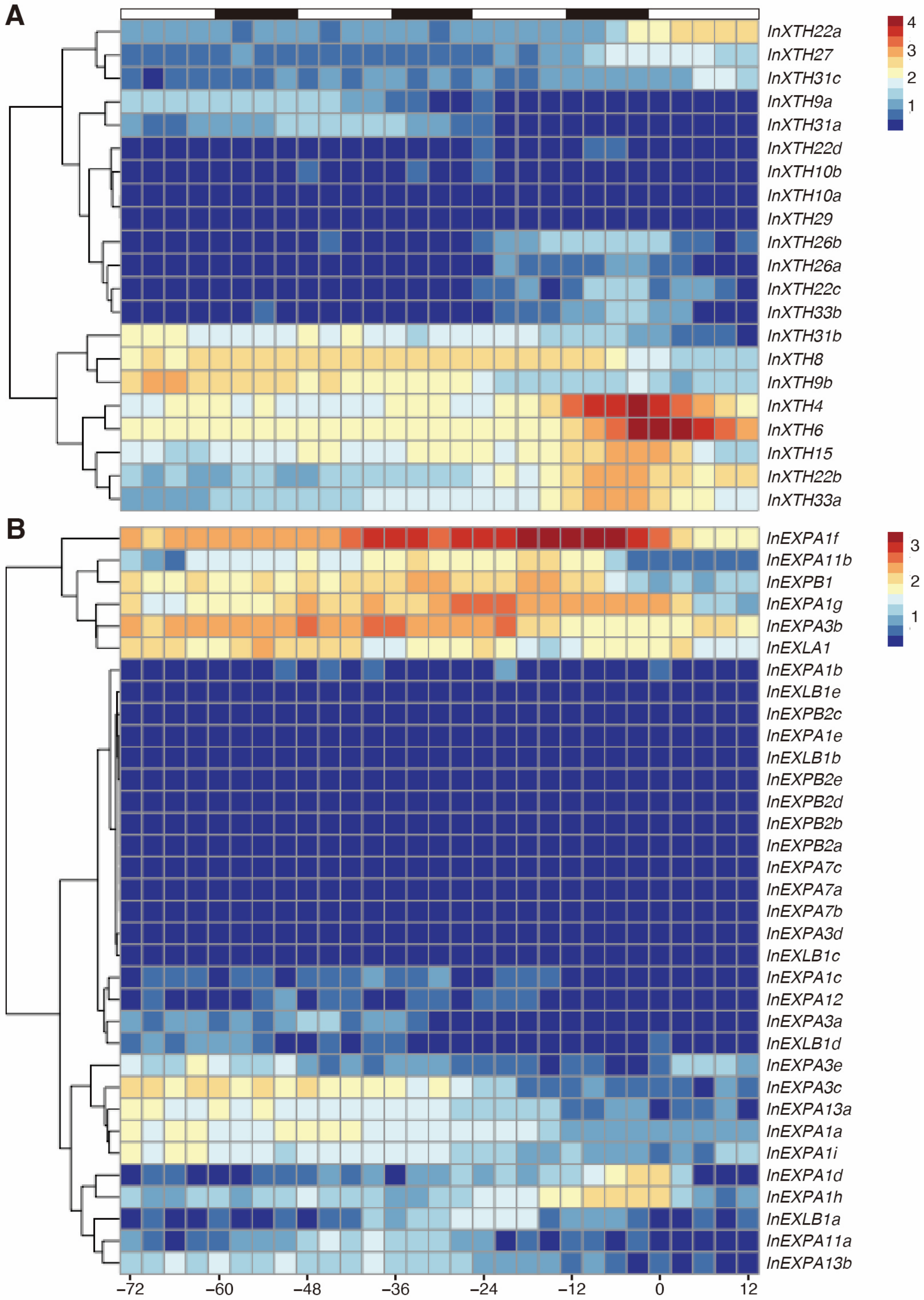


**Figure S2.** Heatmap analysis of the temporal expression patterns of genes involved in cell wall remodeling

Heatmaps showing the expression patterns of *xyloglucan endotransglucosylase/hydrolase* genes (**A**) and *expansin* genes (**B**). The gene lists are provided in Tables S5 and S6, respectively. Expression levels are shown as log10(FPKM + 1). The x-axis indicates time relative to flower opening (0 h). The bars at the top indicate light conditions, with white and black representing light and dark periods, respectively.

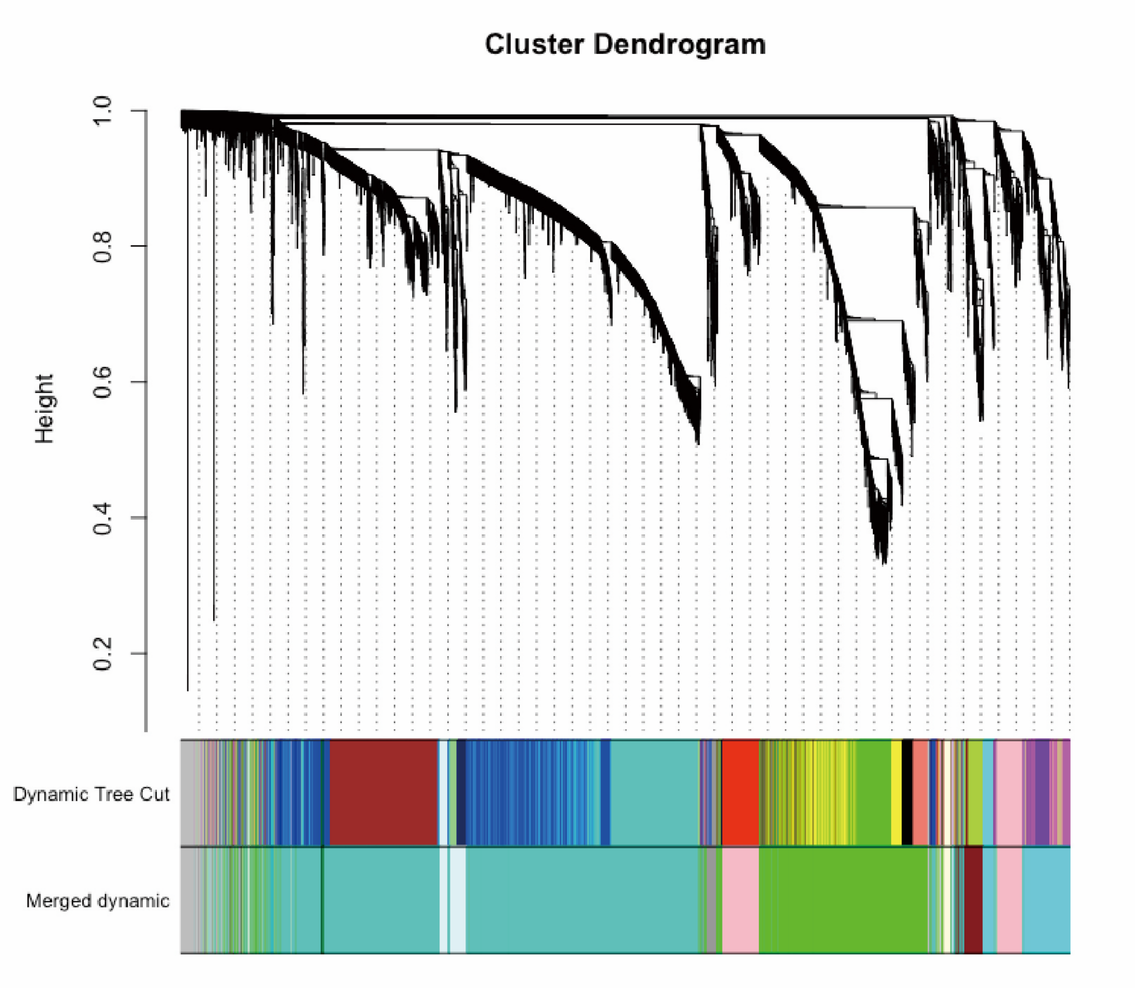


**Figure S3.** Hierarchical clustering analysis by weighted gene co-expression network analysis (WGCNA)

Based on gene expression patterns, 24 co-expression modules were identified by WGCNA and are shown in different colors. Highly correlated modules were subsequently merged, resulting in 11 co-expression modules. The upper panel shows the co-expression modules before merging, and the lower panel shows the merged co-expression modules.

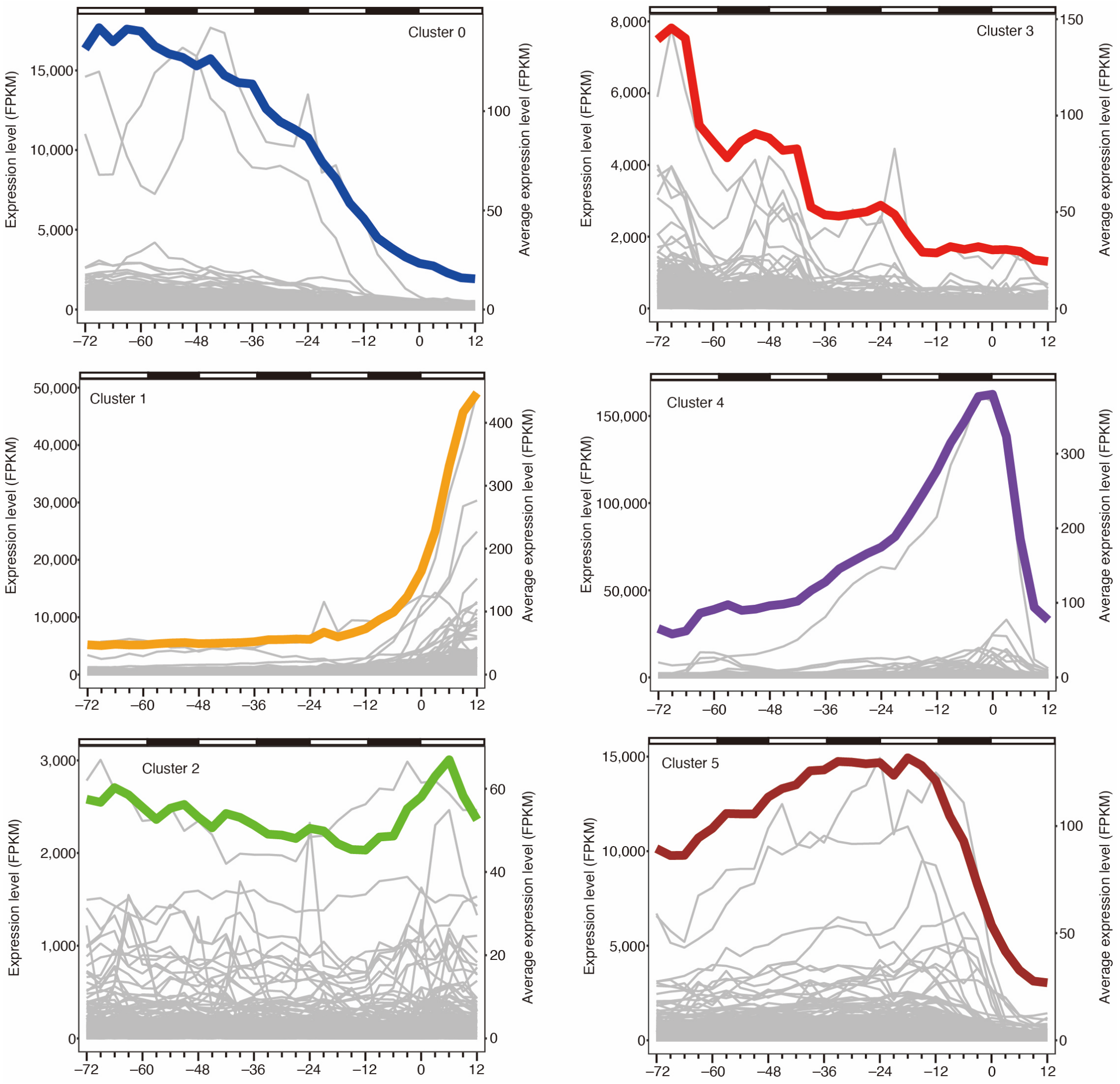


**Figure S4.** Time-series K-means clustering analysis

Changes in gene expression levels for each cluster ($K = 6$). The x-axis indicates time relative to flower opening (0 h). Individual gene expression profiles within each cluster are shown as gray lines using the left y-axis, whereas the mean expression profile of all genes in each cluster is shown as a colored line using the right y-axis.

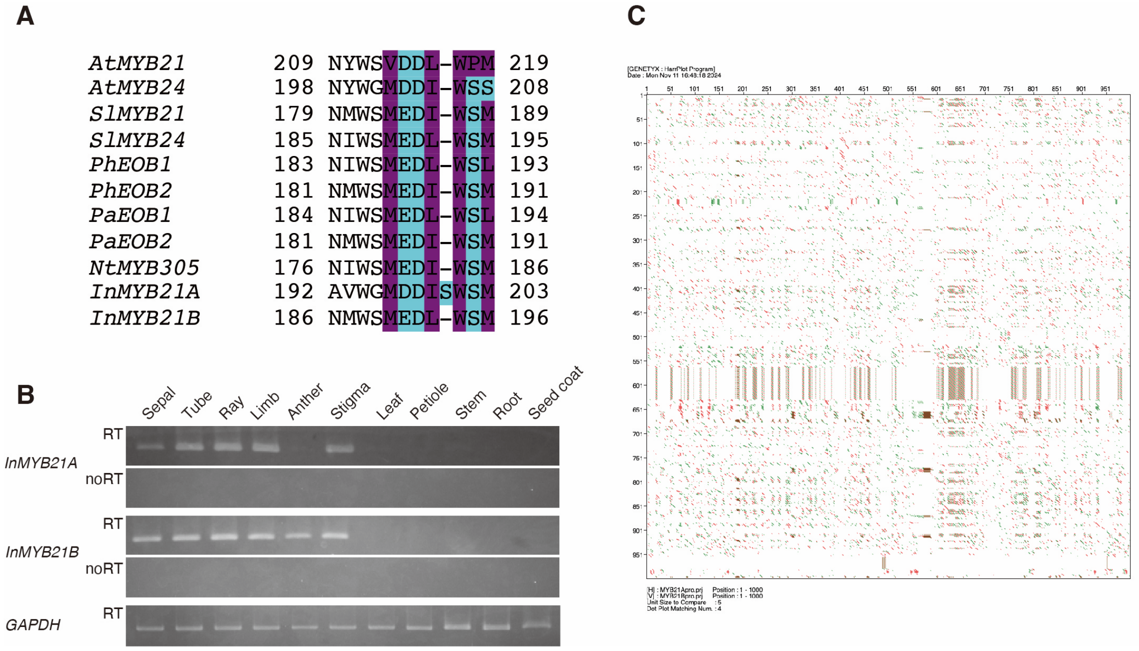


**Figure S5.** Alignment of the transcriptional activation domains of R2R3-MYB subgroup 19 transcription factors, expression analysis of *InMYB21A* and *InMYB21B*, and comparison of their promoter regions

(**A**) Alignment of the amino acid sequences of the transcriptional activation domains of R2R3-MYB subgroup 19 transcription factors. Hydrophilic and hydrophobic amino acids are shown in purple and blue, respectively. (**B**) RT-PCR analysis of *InMYB21A* and *InMYB21B* expression in 11 tissues (sepal, tube, ray, limb, anther, stigma, leaf, petiole, stem, root, and seed coat). Bands from reactions containing reverse transcriptase are labeled RT, whereas those from reactions without reverse transcriptase, used as negative controls, are labeled noRT. The glyceraldehyde-3-phosphate dehydrogenase gene (*GAPDH*) was used as a positive control. (**C**) Harr plots of the 1,000-bp regions upstream of the transcription start sites of *InMYB21A* and *InMYB21B*.

```
Wild type        1630695 AACATGCTGCAATAGTAGCTCTTATGATCCTGAGATCAGGAAGGGACCTTGGACT 1630749
InMYB21A KO #1   1630695 AACATGCTGCAATAGTAGCTCTTATGATCCTGAGATCAGAAGGTAGTTCAGCCAC 1630749
InMYB21A KO #2   1630695 AACATGCTGCAATAGTAGCTCTTATGATCCTGAG--------------------- 1630728
InMYB21A KO #3   1630695 AACATGCTGCAATAGTAGCTCTTATGATCCTGAGAT--GGAAGGGACCTTGGACT 1630747

Wild type        1630750 ATGGAGGAAGACTTGATTCTCATCAACTACATTGCTAATCACGGCGAAGGTGTCT 1630804
InMYB21A KO #1   1630750 CGGAGCCGGCAACTCTTACCGGTACGCTTGAGACCTACATACACAACAAAACAGA 1630804
InMYB21A KO #2   1630729 ----------ACTTGATTCTCATCAACTACATTGCTAATCACGGCGAAGGTGTCT 1630773
InMYB21A KO #3   1630748 ATGGAGGAAGACTTGATTCTCATCAACTACATTGCTAATCACGGCGAAGGTGTCT 1630802

Wild type        1630805 GGAATTCTCTGGCTCGATCCGCAGGTAATTATGATTAAGGAAATACATTAATTTT 1630859
InMYB21A KO #1   1630805 TATTTACTTAATTACATTAATCACTTCCCTAATCTTCAATTGAAAAAAATTAATG 1630859
InMYB21A KO #2   1630774 GGAATTCTCTGGCTCGATCCGCAGGTAATTATGATTAAGGAAATACATTAATTTT 1630828
InMYB21A KO #3   1630803 GGAATTCTCTGGCTCGATCCGCAGGTAATTATGATTAAGGAAATACATTAATTTT 1630857

Wild type        1630860 TTTCAATTGAAGATTAGGGAAGTGATTAATGTAATTAAGTAAATATCTGTTTTGT 1630914
InMYB21A KO #1   1630860 TATTTCCTTAATCATAATTACCTGCGGATCGAGCCAGAGAATTCCAGACACCTTC 1630914
InMYB21A KO #2   1630829 TTTCAATTGAAGATTAGGGAAGTGATTAATGTAATTAAGTAAATATCTGTTTTGT 1630883
InMYB21A KO #3   1630858 TTTCAATTGAAGATTAGGGAAGTGATTAATGTAATTAAGTAAATATCTGTTTTGT 1630912

Wild type        1630915 TGTGTATGTAGGTCTCAAGCGTACCGGTAAGAGTTGCCGGCTCCGGTGGCTGAAC 1630969
InMYB21A KO #1   1630915 GCCGTGATTAGCAATGTAGTTGATGAGAATCAAGTCTTCCTCCATAGTCCAAGGT 1630969
InMYB21A KO #2   1630884 TGTGTATGTAGGTCTCAAGCGTACCGGTAAGAGTTGCCGGCTCCGGTGGCTGAAC 1630938
InMYB21A KO #3   1630913 TGTGTATGTAGGTCTCAAGCGTACCGGTAAGAGTTGCCGGCTCCGGTGGCTGAAC 1630967

Wild type        1630970 TACCTTC-GACCGG 1630982
InMYB21A KO #1   1630970 CCCTTCCTGACCGG 1630983
InMYB21A KO #2   1630939 TACCTTC-GACCGG 1630951
InMYB21A KO #3   1630968 TACCTTC-GACCGG 1630980

Target sequence
PAM sequence
Replaced sequence
```

**Figure S6.** Alleles of *InMYB21A* in genome-edited lines

*InMYB21A* gene sequences in the wild-type and *InMYB21A* genome-edited lines. The guide RNA target sequences are highlighted in yellow, the PAM sequences are highlighted in blue, and replaced sequences are highlighted in gray.

```
Wild type         6487339 TTTTTTTTTTTTGGTTTTTGTAGCAAAACCCTAAAAATTACCAAAATTATGTATT 6487374
InMYB21B KO #1    6487339 TTTTTTTTTTTTGGTTTTTGTAGCAAAACCCTAAAAATTACCAAAATTATGTATT 6487374
InMYB21B KO #2    6487339 TTTTTTTTTTTTGGTTTTTGTATATATAGCACAACATTTCCTAGCTAGCAGTGCA 6487374
InMYB21B KO #3_1  6487339 TTTTTTTTTTTTGGTTTTTGTAGCAAAACCCTAAAAATTACCAAAATTATGTATT 6487374
InMYB21B KO #3_2  6487339 TTTTTTTTTTTTGGTTTTTGTAGCAAAACCCTAAAAATTACCAAAATTATGTATT 6487374
InMYB21B KO #4_1  6487339 TTTTTTTTTTTTGGTTTTTGTAGCAAAACCCTAAAAATTACCAAAATTATGTATT 6487374
InMYB21B KO #4_2  6487339 TTTTTTTTTTTTGGTTTTTGTAGCAAAACCCTAAAAATTACCAAAATTATGTATT 6487374
InMYB21B KO #5_1  6487339 TTTTTTTTTTTTGGTTTTTGTAGCAAAACCCTAAAAATTACCAAAATTATGTATT 6487374
InMYB21B KO #5_2  6487339 TTTTTTTTTTTTGGTTTTTGTAGCAAAACCCTAAAAATTACCAAAATTATGTATT 6487374
InMYB21B KO #6_1  6487339 TTTTTTTTTTTTGGTTTTTGTAGCAAAACCCTAAAAATTACCAAAATTATGTATT 6487374
InMYB21B KO #6_2  6487339 TTTTTTTTTTTTGGTTTTTGTAGCAAAACCCTAAAAATTACCAAAATTATGTATT 6487374

Wild type         6487375 CACTCCATGTTTTGATTTTGTTTTGTTGGTATAGGTCTGAAGCGTACCGGGAAGA 6487429
InMYB21B KO #1    6487375 CACTCCATGTTTTGATTTTGTTTTGTTGGTATAGGTCTGAAGCGTACCGGGAAGA 6487429
InMYB21B KO #2    6487375 GTTGGAATCATGATGAAAAGGAAGCTCCAACTTGCTTTTACGGCGAATAATCGAA 6487429
InMYB21B KO #3_1  6487375 CACTCCATGTTTTGATTTTGTTTTGTTGGTATAGGTCTGAAGCGTACCGGGAA-- 6487427
InMYB21B KO #3_2  6487375 CACTCCATGTTTTGATTTTGTTTTGTTGGTATAGGTCTGAAGCGTACCGGGAAGA 6487429
InMYB21B KO #4_1  6487375 CACTCCATGTTTTGATTTTGTTTTGTTGGTATAGGTCTGAAGCGTACCGGGAAGA 6487429
InMYB21B KO #4_2  6487375 CACTCCATGTTTTGATTTTGTTTTGTTGGTATAGGTCTGAAGCGTACCGGGAAGA 6487429
InMYB21B KO #5_1  6487375 CACTCCATGTTTTGATTTTGTTTTGTTGGTATAGGTCTGAAGCGTACCGGGAAGA 6487429
InMYB21B KO #5_2  6487375 CACTCCATGTTTTGATTTTGTTTTGTTGGTATAGGTCTGAAGCGTACCGGGAAGA 6487429
InMYB21B KO #6_1  6487375 CACTCCATGTTTTGATTTTGTTTTGTTGGTATAGGTCTGAAGCGTACCGGGAAGA 6487429
InMYB21B KO #6_2  6487375 CACTCCATGTTTTGATTTTGTTTTGTTGGTATAGGTCTGAAGCGTACCGGGAAGA 6487429

Wild type         6487430 GTTGCCGGCTCCGGTGGCTAAACTACCTCCGGCCAGAC-GTCCGGCGGGGGAATA 6487483
InMYB21B KO #1    6487430 GTTGCCGGCTCCGGTGGCTAAACTACCTCCGGCCAG----TCCGGCGGGGGAATA 6487480
InMYB21B KO #2    6487430 TTCTTACGGCGAATAATCACGAATTCTTGCTTTTG-------------------- 6487464
InMYB21B KO #3_1  6487427 ------------------------------------------------------- 6487427
InMYB21B KO #3_2  6487430 GTTGCCGGCTCCGGTGGCTAAACTACCTCCGG-------------CGGGGGAATA 6487471
InMYB21B KO #4_1  6487430 GTTGCCGGCTCCGGTGGCTAAACTACCTCCGGA-------------GGGGGAATA 6487471
InMYB21B KO #4_2  6487430 GTTGCCGGCTCCGGTGGCTAAACTACCTCCGGCCAGACCGTCCGGCGGGGGAATA 6487484
InMYB21B KO #5_1  6487430 GTTGCCGGCTCCGGTGGCTAAACTACCTCCGGCCAGAC-G--------------- 6487468
InMYB21B KO #5_2  6487430 GTTGCCGGCTCCGGTGGCTAAACTACCTCCGGCC-----GTCCGGCGGGGGAATA 6487479
InMYB21B KO #6_1  6487430 GTTGCCGGCTCCGGTGGCTAAACTACCTCCGGCCAGAC-G--------------- 6487468
InMYB21B KO #6_2  6487430 GTTGCCGGCTCCGGTGGCTAAACTACCTCCGGCC---C-GTCCGGCGGGGGAATA 6487480

Wild type         6487484 TTACGCCGGAGG-AACAACTCCTGATTATGGAATTGCATGCTAAGTGGGAAATA 6487537
InMYB21B KO #1    6487481 TTACGCCGGAGGGAACAACTCCTGATTATGGAATTGCATGCTAAGTGGGAAATA 6487534
InMYB21B KO #2    6487464 ------------------------------------------------------ 6487464
InMYB21B KO #3_1  6487427 -----------G-AACAACTCCTGATTATGGAATTGCATGCTAAGTGGGAAATA 6487469
InMYB21B KO #3_2  6487472 TTACGCCGGAGG-AACAACTCCTGATTATGGAATTGCATGCTAAGTGGGAAATA 6487526
InMYB21B KO #4_1  6487472 TTACGCCGGAGGTAACAACTCCTGATTATGGAATTGCATGCTAAGTGGGAAATA 6487527
InMYB21B KO #4_2  6487485 TTACGCCGGAG----CAACTCCTGATTATGGAATTGCATGCTAAGTGGGAAATA 6487535
InMYB21B KO #5_1  6487469 ------------AACAACTCCTGATTATGGAATTGCATGCTAAGTGGGAAATA 6487510
InMYB21B KO #5_2  6487480 TTACGCCGGAGGTAACAACTCCTGATTATGGAATTGCATGCTAAGTGGGAAATA 6487535
InMYB21B KO #6_1  6487469 ------------AACAACTCCTGATTATGGAATTGCATGCTAAGTGGGAAATA 6487510
InMYB21B KO #6_2  6487481 TTACGCCGGAG--AACAACTCCTGATTATGGAATTGCATGCTAAGTGGGAAATA 6487533

Wild type         6487538 GGTGAGTGAGTATATATATTCTTTAATTTTGAGTTTTATATATGTTGATGAATTG 6487592
InMYB21B KO #1    6487535 GGTGAGTGAGTATATATATTCTTTAATTTTGAGTTTTATATATGTTGATGAATTG 6487589
InMYB21B KO #2    6487464 ------------------------------------------------------- 6487464
InMYB21B KO #3_1  6487470 GGTGAGTGAGTATATATATTCTTTAATTTTGAGTTTTATATATGTTGATGAATTG 6487524
InMYB21B KO #3_2  6487527 GGTGAGTGAGTATATATATTCTTTAATTTTGAGTTTTATATATGTTGATGAATTG 6487581
InMYB21B KO #4_1  6487528 GGTGAGTGAGTATATATATTCTTTAATTTTGAGTTTTATATATGTTGATGAATTG 6487582
InMYB21B KO #4_2  6487536 GGTGAGTGAGTATATATATTCTTTAATTTTGAGTTTTATATATGTTGATGAATTG 6487590
InMYB21B KO #5_1  6487511 GGTGAGTGAGTATATATATTCTTTAATTTTGAGTTTTATATATGTTGATGAATTG 6487565
InMYB21B KO #5_2  6487536 GGTGAGTGAGTATATATATTCTTTAATTTTGAGTTTTATATATGTTGATGAATTG 6487590
InMYB21B KO #6_1  6487511 GGTGAGTGAGTATATATATTCTTTAATTTTGAGTTTTATATATGTTGATGAATTG 6487565
InMYB21B KO #6_2  6487534 GGTGAGTGAGTATATATATTCTTTAATTTTGAGTTTTATATATGTTGATGAATTG 6487588

Wild type         6487593 CTAGAACAACAAATTTTAATTTGTATATATATAAAAAAAAATGTGGAATTTTCTC 6487647
InMYB21B KO #1    6487590 CTAGAACAACAAATTTTAATTTGTATATATATAAAAAAAAATGTGGAATTTTCTC 6487644
InMYB21B KO #2    6487464 ------------------------------------------------------- 6487464
InMYB21B KO #3_1  6487525 CTAGAACAACAAATTTTAATTTGTATATATATAAAAAAAAATGTGGAATTTTCTC 6487579
InMYB21B KO #3_2  6487582 CTAGAACAACAAATTTTAATTTGTATATATATAAAAAAAAATGTGGAATTTTCTC 6487636
InMYB21B KO #4_1  6487583 CTAGAACAACAAATTTTAATTTGTATATATATAAAAAAAAATGTGGAATTTTCTC 6487637
InMYB21B KO #4_2  6487591 CTAGAACAACAAATTTTAATTTGTATATATATAAAAAAAAATGTGGAATTTTCTC 6487645
InMYB21B KO #5_1  6487567 CTAGAACAACAAATTTTAATTTGTATATATATAAAAAAAAATGTGGAATTTTCTC 6487620
InMYB21B KO #5_2  6487591 CTAGAACAACAAATTTTAATTTGTATATATATAAAAAAAAATGTGGAATTTTCTC 6487645
InMYB21B KO #6_1  6487567 CTAGAACAACAAATTTTAATTTGTATATATATAAAAAAAAATGTGGAATTTTCTC 6487620
InMYB21B KO #6_2  6487589 CTAGAACAACAAATTTTAATTTGTATATATATAAAAAAAAATGTGGAATTTTCTC 6487643

Wild type         6487648 ATCCAAGAATTCGATTATTCGCCGTAAGACAAAGAAGAGATGAGCCGCAAGCTAT 6487702
InMYB21B KO #1    6487645 ATCCAAGAATTCGATTATTCGCCGTAAGACAAAGAAGAGATGAGCCGCAAGCTAT 6487699
InMYB21B KO #2    6487464 ------------------------------------------------------- 6487464
InMYB21B KO #3_1  6487580 ATCCAAGAATTCGATTATTCGCCGTAAGACAAAGAAGAGATGAGCCGCAAGCTAT 6487634
InMYB21B KO #3_2  6487637 ATCCAAGAATTCGATTATTCGCCGTAAGACAAAGAAGAGATGAGCCGCAAGCTAT 6487691
InMYB21B KO #4_1  6487638 ATCCAAGAATTCGATTATTCGCCGTAAGACAAAGAAGAGATGAGCCGCAAGCTAT 6487692
InMYB21B KO #4_2  6487646 ATCCAAGAATTCGATTATTCGCCGTAAGACAAAGAAGAGATGAGCCGCAAGCTAT 6487700
InMYB21B KO #5_1  6487621 ATCCAAGAATTCGATTATTCGCCGTAAGACAAAGAAGAGATGAGCCGCAAGCTAT 6487675
InMYB21B KO #5_2  6487646 ATCCAAGAATTCGATTATTCGCCGTAAGACAAAGAAGAGATGAGCCGCAAGCTAT 6487700
InMYB21B KO #6_1  6487621 ATCCAAGAATTCGATTATTCGCCGTAAGACAAAGAAGAGATGAGCCGCAAGCTAT 6487675
```

*InMYB21B* KO #6_2  6487644 ATCCAAGAATTCGATTATTCGCCGTAAGACAAAGAAGAGATGAGCCGCAAGCTAT 6487698

Wild type           6487703 ATGTGTTATGTGTACCCTTTTGTGTCGCCCAAAAAAATAAATAAATAAAGAATCAA 6487757
*InMYB21B* KO #1    6487700 ATGTGTTATGTGTACCCTTTTGTGTCGCCCAAAAAAATAAATAAATAAAGAATCAA 6487754
*InMYB21B* KO #2    6487464 ------------------------------------------------------- 6487464
*InMYB21B* KO #3_1  6487635 ATGTGTTATGTGTACCCTTTTGTGTCGCCCAAAAAAATAAATAAATAAAGAATCAA 6487689
*InMYB21B* KO #3_2  6487692 ATGTGTTATGTGTACCCTTTTGTGTCGCCCAAAAAAATAAATAAATAAAGAATCAA 6487746
*InMYB21B* KO #4_1  6487693 ATGTGTTATGTGTACCCTTTTGTGTCGCCCAAAAAAATAAATAAATAAAGAATCAA 6487747
*InMYB21B* KO #4_2  6487701 ATGTGTTATGTGTACCCTTTTGTGTCGCCCAAAAAAATAAATAAATAAAGAATCAA 6487755
*InMYB21B* KO #5_1  6487676 ATGTGTTATGTGTACCCTTTTGTGTCGCCCAAAAAAATAAATAAATAAAGAATCAA 6487730
*InMYB21B* KO #5_2  6487701 ATGTGTTATGTGTACCCTTTTGTGTCGCCCAAAAAAATAAATAAATAAAGAATCAA 6487755
*InMYB21B* KO #6_1  6487676 ATGTGTTATGTGTACCCTTTTGTGTCGCCCAAAAAAATAAATAAATAAAGAATCAA 6487730
*InMYB21B* KO #6_2  6487699 ATGTGTTATGTGTACCCTTTTGTGTCGCCCAAAAAAATAAATAAATAAAGAATCAA 6487753

Wild type           6487758 AATCCATAGCTTGTGGCTGTGTGATCTCTTCAAGTCAAAGTGAGTTATTAGTTCA 6487812
*InMYB21B* KO #1    6487755 AATCCATAGCTTGTGGCTGTGTGATCTCTTCAAGTCAAAGTGAGTTATTAGTTCA 6487809
*InMYB21B* KO #2    6487464 ------------------------------------------------------- 6487464
*InMYB21B* KO #3_1  6487690 AATCCATAGCTTGTGGCTGTGTGATCTCTTCAAGTCAAAGTGAGTTATTAGTTCA 6487744
*InMYB21B* KO #3_2  6487747 AATCCATAGCTTGTGGCTGTGTGATCTCTTCAAGTCAAAGTGAGTTATTAGTTCA 6487801
*InMYB21B* KO #4_1  6487748 AATCCATAGCTTGTGGCTGTGTGATCTCTTCAAGTCAAAGTGAGTTATTAGTTCA 6487802
*InMYB21B* KO #4_2  6487756 AATCCATAGCTTGTGGCTGTGTGATCTCTTCAAGTCAAAGTGAGTTATTAGTTCA 6487810
*InMYB21B* KO #5_1  6487731 AATCCATAGCTTGTGGCTGTGTGATCTCTTCAAGTCAAAGTGAGTTATTAGTTCA 6487785
*InMYB21B* KO #5_2  6487756 AATCCATAGCTTGTGGCTGTGTGATCTCTTCAAGTCAAAGTGAGTTATTAGTTCA 6487810
*InMYB21B* KO #6_1  6487731 AATCCATAGCTTGTGGCTGTGTGATCTCTTCAAGTCAAAGTGAGTTATTAGTTCA 6487785
*InMYB21B* KO #6_2  6487754 AATCCATAGCTTGTGGCTGTGTGATCTCTTCAAGTCAAAGTGAGTTATTAGTTCA 6487808

Wild type           6487813 TTGGTAGAGAGAATAGCAACGTGGGGATTATGATCTCTTCAATCCACGTGCGGAA 6487868
*InMYB21B* KO #1    6487810 TTGGTAGAGAGAATAGCAACGTGGGGATTATGATCTCTTCAATCCACGTGCGGAA 6487865
*InMYB21B* KO #2    6487464 ------------------------------------------------------- 6487464
*InMYB21B* KO #3_1  6487745 TTGGTAGAGAGAATAGCAACGTGGGGATTATGATCTCTTCAATCCACGTGCGGAA 6487799
*InMYB21B* KO #3_2  6487802 TTGGTAGAGAGAATAGCAACGTGGGGATTATGATCTCTTCAATCCACGTGCGGAA 6487856
*InMYB21B* KO #4_1  6487803 TTGGTAGAGAGAATAGCAACGTGGGGATTATGATCTCTTCAATCCACGTGCGGAA 6487857
*InMYB21B* KO #4_2  6487811 TTGGTAGAGAGAATAGCAACGTGGGGATTATGATCTCTTCAATCCACGTGCGGAA 6487866
*InMYB21B* KO #5_1  6487786 TTGGTAGAGAGAATAGCAACGTGGGGATTATGATCTCTTCAATCCACGTGCGGAA 6487840
*InMYB21B* KO #5_2  6487811 TTGGTAGAGAGAATAGCAACGTGGGGATTATGATCTCTTCAATCCACGTGCGGAA 6487866
*InMYB21B* KO #6_1  6487786 TTGGTAGAGAGAATAGCAACGTGGGGATTATGATCTCTTCAATCCACGTGCGGAA 6487840
*InMYB21B* KO #6_2  6487809 TTGGTAGAGAGAATAGCAACGTGGGGATTATGATCTCTTCAATCCACGTGCGGAA 6487863

Wild type           6487869 ATCAATATATGGTCAGTGTTCATCCTACTAAGATTTCTGCCTAGCTTTTCCCAGA 6487922
*InMYB21B* KO #1    6487866 ATCAATATATGGTCAGTGTTCATCCTACTAAGATTTCTGCCTAGCTTTTCCCAGA 6487919
*InMYB21B* KO #2    6487464 ------------------------------------------------------- 6487464
*InMYB21B* KO #3_1  6487800 ATCAATATATGGTCAGTGTTCATCCTACTAAGATTTCTGCCTAGCTTTTCCCAGA 6487854
*InMYB21B* KO #3_2  6487857 ATCAATATATGGTCAGTGTTCATCCTACTAAGATTTCTGCCTAGCTTTTCCCAGA 6487911
*InMYB21B* KO #4_1  6487858 ATCAATATATGGTCAGTGTTCATCCTACTAAGATTTCTGCCTAGCTTTTCCCAGA 6487912
*InMYB21B* KO #4_2  6487867 ATCAATATATGGTCAGTGTTCATCCTACTAAGATTTCTGCCTAGCTTTTCCCAGA 6487920
*InMYB21B* KO #5_1  6487841 ATCAATATATGGTCAGTGTTCATCCTACTAAGATTTCTGCCTAGCTTTTCCCAGA 6487895
*InMYB21B* KO #5_2  6487867 ATCAATATATGGTCAGTGTTCATCCTACTAAGATTTCTGCCTAGCTTTTCCCAGA 6487920
*InMYB21B* KO #6_1  6487841 ATCAATATATGGTCAGTGTTCATCCTACTAAGATTTCTGCCTAGCTTTTCCCAGA 6487895
*InMYB21B* KO #6_2  6487864 ATCAATATATGGTCAGTGTTCATCCTACTAAGATTTCTGCCTAGCTTTTCCCAGA 6487918

Wild type           6487923 TTTAATTTGTTCTCAAACTTTTTCCCAAAAAAATAAAGAAAAAAAAAAACTCAAGAA 6487977
*InMYB21B* KO #1    6487920 TTTAATTTGTTCTCAAACTTTTTCCCAAAAAAATAAAGAAAAAAAAAAACTCAAGAA 6487974
*InMYB21B* KO #2    6487464 ------------------------------------------------------- 6487464
*InMYB21B* KO #3_1  6487855 TTTAATTTGTTCTCAAACTTTTTCCCAAAAAAATAAAGAAAAAAAAAAACTCAAGAA 6487909
*InMYB21B* KO #3_2  6487912 TTTAATTTGTTCTCAAACTTTTTCCCAAAAAAATAAAGAAAAAAAAAAACTCAAGAA 6487966
*InMYB21B* KO #4_1  6487913 TTTAATTTGTTCTCAAACTTTTTCCCAAAAAAATAAAGAAAAAAAAAAACTCAAGAA 6487967
*InMYB21B* KO #4_2  6487921 TTTAATTTGTTCTCAAACTTTTTCCCAAAAAAATAAAGAAAAAAAAAAACTCAAGAA 6487975
*InMYB21B* KO #5_1  6487896 TTTAATTTGTTCTCAAACTTTTTCCCAAAAAAATAAAGAAAAAAAAAAACTCAAGAA 6487950
*InMYB21B* KO #5_2  6487921 TTTAATTTGTTCTCAAACTTTTTCCCAAAAAAATAAAGAAAAAAAAAAACTCAAGAA 6487975
*InMYB21B* KO #6_1  6487896 TTTAATTTGTTCTCAAACTTTTTCCCAAAAAAATAAAGAAAAAAAAAAACTCAAGAA 6487950
*InMYB21B* KO #6_2  6487919 TTTAATTTGTTCTCAAACTTTTTCCCAAAAAAATAAAGAAAAAAAAAAACTCAAGAA 6487973

Wild type           6487978 ATTCTACTTGCTACCTTGTTTTTACTTAATTATTAGTTTCTTTGATGAAGTTCAT 6488032
*InMYB21B* KO #1    6487975 ATTCTACTTGCTACCTTGTTTTTACTTAATTATTAGTTTCTTTGATGAAGTTCAT 6488029
*InMYB21B* KO #2    6487464 ------------------------------------------------------- 6487464
*InMYB21B* KO #3_1  6487910 ATTCTACTTGCTACCTTGTTTTTACTTAATTATTAGTTTCTTTGATGAAGTTCAT 6487964
*InMYB21B* KO #3_2  6487967 ATTCTACTTGCTACCTTGTTTTTACTTAATTATTAGTTTCTTTGATGAAGTTCAT 6488021
*InMYB21B* KO #4_1  6487968 ATTCTACTTGCTACCTTGTTTTTACTTAATTATTAGTTTCTTTGATGAAGTTCAT 6488022
*InMYB21B* KO #4_2  6487976 ATTCTACTTGCTACCTTGTTTTTACTTAATTATTAGTTTCTTTGATGAAGTTCAT 6488030
*InMYB21B* KO #5_1  6487951 ATTCTACTTGCTACCTTGTTTTTACTTAATTATTAGTTTCTTTGATGAAGTTCAT 6488005
*InMYB21B* KO #5_2  6487976 ATTCTACTTGCTACCTTGTTTTTACTTAATTATTAGTTTCTTTGATGAAGTTCAT 6488030
*InMYB21B* KO #6_1  6487951 ATTCTACTTGCTACCTTGTTTTTACTTAATTATTAGTTTCTTTGATGAAGTTCAT 6488005
*InMYB21B* KO #6_2  6487974 ATTCTACTTGCTACCTTGTTTTTACTTAATTATTAGTTTCTTTGATGAAGTTCAT 6488028

Wild type           6488033 AAAATTAGCATATAACATCATTATTTGATTGCTTTTTTTAAAAAAAAAAATATGTA 6488087
*InMYB21B* KO #1    6488030 AAAATTAGCATATAACATCATTATTTGATTGCTTTTTTTAAAAAAAAAAATATGTA 6488084
*InMYB21B* KO #2    6487464 ------------------------------------------------------- 6487464
*InMYB21B* KO #3_1  6487965 AAAATTAGCATATAACATCATTATTTGATTGCTTTTTTTAAAAAAAAAAATATGTA 6488019
*InMYB21B* KO #3_2  6488022 AAAATTAGCATATAACATCATTATTTGATTGCTTTTTTTAAAAAAAAAAATATGTA 6488076
*InMYB21B* KO #4_1  6488023 AAAATTAGCATATAACATCATTATTTGATTGCTTTTTTTAAAAAAAAAAATATGTA 6488077
*InMYB21B* KO #4_2  6488031 AAAATTAGCATATAACATCATTATTTGATTGCTTTTTTTAAAAAAAAAAATATGTA 6488085
*InMYB21B* KO #5_1  6488006 AAAATTAGCATATAACATCATTATTTGATTGCTTTTTTTAAAAAAAAAAATATGTA 6488060

*InMYB21B* KO #5_2 6488031 AAAATTAGCATATAACATCATTATTTGATTGCTTTTTTTAAAAAAAAAAATATGTA 6488085
*InMYB21B* KO #6_1 6488006 AAAATTAGCATATAACATCATTATTTGATTGCTTTTTTTAAAAAAAAAAATATGTA 6488060
*InMYB21B* KO #6_2 6488029 AAAATTAGCATATAACATCATTATTTGATTGCTTTTTTTAAAAAAAAAAATATGTA 6488083

Wild type 6488088 AAATGATTAATGAATCCCCTAATTGAAATATAATGTTAGATTCTATTCTTTTAAA 6488142
*InMYB21B* KO #1 6488085 AAATGATTAATGAATCCCCTAATTGAAATATAATGTTAGATTCTATTCTTTTAAA 6488139
*InMYB21B* KO #2 6487464 ------------------------------------------------------- 6487464
*InMYB21B* KO #3_1 6488020 AAATGATTAATGAATCCCCTAATTGAAATATAATGTTAGATTCTATTCTTTTAAA 6488074
*InMYB21B* KO #3_2 6488077 AAATGATTAATGAATCCCCTAATTGAAATATAATGTTAGATTCTATTCTTTTAAA 6488131
*InMYB21B* KO #4_1 6488078 AAATGATTAATGAATCCCCTAATTGAAATATAATGTTAGATTCTATTCTTTTAAA 6488132
*InMYB21B* KO #4_2 6488086 AAATGATTAATGAATCCCCTAATTGAAATATAATGTTAGATTCTATTCTTTTAAA 6488140
*InMYB21B* KO #5_1 6488061 AAATGATTAATGAATCCCCTAATTGAAATATAATGTTAGATTCTATTCTTTTAAA 6488115
*InMYB21B* KO #5_2 6488086 AAATGATTAATGAATCCCCTAATTGAAATATAATGTTAGATTCTATTCTTTTAAA 6488140
*InMYB21B* KO #6_1 6488061 AAATGATTAATGAATCCCCTAATTGAAATATAATGTTAGATTCTATTCTTTTAAA 6488115
*InMYB21B* KO #6_2 6488084 AAATGATTAATGAATCCCCTAATTGAAATATAATGTTAGATTCTATTCTTTTAAA 6488138

Wild type 6488143 TTAATAAATTAATTATTCTTAACTGATTGCTTAAACATGATTAATTAGCATGTAA 6488197
*InMYB21B* KO #1 6488140 TTAATAAATTAATTATTCTTAACTGATTGCTTAAACATGATTAATTAGCATGTAA 6488194
*InMYB21B* KO #2 6487464 ------------------------------------------------------- 6487464
*InMYB21B* KO #3_1 6488075 TTAATAAATTAATTATTCTTAACTGATTGCTTAAACATGATTAATTAGCATGTAA 6488129
*InMYB21B* KO #3_2 6488132 TTAATAAATTAATTATTCTTAACTGATTGCTTAAACATGATTAATTAGCATGTAA 6488186
*InMYB21B* KO #4_1 6488133 TTAATAAATTAATTATTCTTAACTGATTGCTTAAACATGATTAATTAGCATGTAA 6488187
*InMYB21B* KO #4_2 6488141 TTAATAAATTAATTATTCTTAACTGATTGCTTAAACATGATTAATTAGCATGTAA 6488195
*InMYB21B* KO #5_1 6488116 TTAATAAATTAATTATTCTTAACTGATTGCTTAAACATGATTAATTAGCATGTAA 6488170
*InMYB21B* KO #5_2 6488141 TTAATAAATTAATTATTCTTAACTGATTGCTTAAACATGATTAATTAGCATGTAA 6488195
*InMYB21B* KO #6_1 6488116 TTAATAAATTAATTATTCTTAACTGATTGCTTAAACATGATTAATTAGCATGTAA 6488170
*InMYB21B* KO #6_2 6488139 TTAATAAATTAATTATTCTTAACTGATTGCTTAAACATGATTAATTAGCATGTAA 6488193

Wild type 6488198 TCACTAACTTATCGTGCTCACAGTGAGTATATATCTATATATATTGTAATCACTA 6488252
*InMYB21B* KO #1 6488195 TCACTAACTTATCGTGCTCACAGTGAGTATATATCTATATATATTGTAATCACTA 6488249
*InMYB21B* KO #2 6487464 ------------------------------------------------------- 6487464
*InMYB21B* KO #3_1 6488130 TCACTAACTTATCGTGCTCACAGTGAGTATATATCTATATATATTGTAATCACTA 6488184
*InMYB21B* KO #3_2 6488187 TCACTAACTTATCGTGCTCACAGTGAGTATATATCTATATATATTGTAATCACTA 6488241
*InMYB21B* KO #4_1 6488188 TCACTAACTTATCGTGCTCACAGTGAGTATATATCTATATATATTGTAATCACTA 6488242
*InMYB21B* KO #4_2 6488196 TCACTAACTTATCGTGCTCACAGTGAGTATATATCTATATATATTGTAATCACTA 6488250
*InMYB21B* KO #5_1 6488171 TCACTAACTTATCGTGCTCACAGTGAGTATATATCTATATATATTGTAATCACTA 6488225
*InMYB21B* KO #5_2 6488196 TCACTAACTTATCGTGCTCACAGTGAGTATATATCTATATATATTGTAATCACTA 6488250
*InMYB21B* KO #6_1 6488171 TCACTAACTTATCGTGCTCACAGTGAGTATATATCTATATATATTGTAATCACTA 6488225
*InMYB21B* KO #6_2 6488194 TCACTAACTTATCGTGCTCACAGTGAGTATATATCTATATATATTGTAATCACTA 6488248

Wild type 6488253 ACTTATCGTGCCCACAGTGAGTATATATCTATATATATTATAATATTTTATATCA 6488307
*InMYB21B* KO #1 6488250 ACTTATCGTGCCCACAGTGAGTATATATCTATATATATTATAATATTTTATATCA 6488304
*InMYB21B* KO #2 6487464 ------------------------------------------------------- 6487464
*InMYB21B* KO #3_1 6488185 ACTTATCGTGCCCACAGTGAGTATATATCTATATATATTATAATATTTTATATCA 6488239
*InMYB21B* KO #3_2 6488242 ACTTATCGTGCCCACAGTGAGTATATATCTATATATATTATAATATTTTATATCA 6488296
*InMYB21B* KO #4_1 6488243 ACTTATCGTGCCCACAGTGAGTATATATCTATATATATTATAATATTTTATATCA 6488297
*InMYB21B* KO #4_2 6488251 ACTTATCGTGCCCACAGTGAGTATATATCTATATATATTATAATATTTTATATCA 6488305
*InMYB21B* KO #5_1 6488226 ACTTATCGTGCCCACAGTGAGTATATATCTATATATATTATAATATTTTATATCA 6488280
*InMYB21B* KO #5_2 6488251 ACTTATCGTGCCCACAGTGAGTATATATCTATATATATTATAATATTTTATATCA 6488305
*InMYB21B* KO #6_1 6488226 ACTTATCGTGCCCACAGTGAGTATATATCTATATATATTATAATATTTTATATCA 6488280
*InMYB21B* KO #6_2 6488249 ACTTATCGTGCCCACAGTGAGTATATATCTATATATATTATAATATTTTATATCA 6488303

Wild type 6488308 TAAAAAGTACTCGGTGACGAATAAATTAATAAAAAAGGGTATTGTAGAAATAGGG 6488362
*InMYB21B* KO #1 6488305 TAAAAAGTACTCGGTGACGAATAAATTAATAAAAAAGGGTATTGTAGAAATAGGG 6488359
*InMYB21B* KO #2 6487464 ------------------------------------------------------- 6487464
*InMYB21B* KO #3_1 6488240 TAAAAAGTACTCGGTGACGAATAAATTAATAAAAAAGGGTATTGTAGAAATAGGG 6488294
*InMYB21B* KO #3_2 6488297 TAAAAAGTACTCGGTGACGAATAAATTAATAAAAAAGGGTATTGTAGAAATAGGG 6488351
*InMYB21B* KO #4_1 6488298 TAAAAAGTACTCGGTGACGAATAAATTAATAAAAAAGGGTATTGTAGAAATAGGG 6488352
*InMYB21B* KO #4_2 6488306 TAAAAAGTACTCGGTGACGAATAAATTAATAAAAAAGGGTATTGTAGAAATAGGG 6488360
*InMYB21B* KO #5_1 6488281 TAAAAAGTACTCGGTGACGAATAAATTAATAAAAAAGGGTATTGTAGAAATAGGG 6488335
*InMYB21B* KO #5_2 6488306 TAAAAAGTACTCGGTGACGAATAAATTAATAAAAAAGGGTATTGTAGAAATAGGG 6488360
*InMYB21B* KO #6_1 6488281 TAAAAAGTACTCGGTGACGAATAAATTAATAAAAAAGGGTATTGTAGAAATAGGG 6488335
*InMYB21B* KO #6_2 6488304 TAAAAAGTACTCGGTGACGAATAAATTAATAAAAAAGGGTATTGTAGAAATAGGG 6488358

Wild type 6488363 AGATGAAGTATTTTATATATATTTTTTTGTTTTTAAAAGAAGTGATTTTAAGGTGG 6488417
*InMYB21B* KO #1 6488360 AGATGAAGTATTTTATATATATTTTTTTGTTTTTAAAAGAAGTGATTTTAAGGTGG 6488414
*InMYB21B* KO #2 6487464 ------------------------------------------------------- 6487464
*InMYB21B* KO #3_1 6488295 AGATGAAGTATTTTATATATATTTTTTTGTTTTTAAAAGAAGTGATTTTAAGGTGG 6488349
*InMYB21B* KO #3_2 6488352 AGATGAAGTATTTTATATATATTTTTTTGTTTTTAAAAGAAGTGATTTTAAGGTGG 6488406
*InMYB21B* KO #4_1 6488353 AGATGAAGTATTTTATATATATTTTTTTGTTTTTAAAAGAAGTGATTTTAAGGTGG 6488407
*InMYB21B* KO #4_2 6488361 AGATGAAGTATTTTATATATATTTTTTTGTTTTTAAAAGAAGTGATTTTAAGGTGG 6488415
*InMYB21B* KO #5_1 6488336 AGATGAAGTATTTTATATATATTTTTTTGTTTTTAAAAGAAGTGATTTTAAGGTGG 6488390
*InMYB21B* KO #5_2 6488361 AGATGAAGTATTTTATATATATTTTTTTGTTTTTAAAAGAAGTGATTTTAAGGTGG 6488415
*InMYB21B* KO #6_1 6488336 AGATGAAGTATTTTATATATATTTTTTTGTTTTTAAAAGAAGTGATTTTAAGGTGG 6488390
*InMYB21B* KO #6_2 6488359 AGATGAAGTATTTTATATATATTTTTTTGTTTTTAAAAGAAGTGATTTTAAGGTGG 6488413

Wild type 6488418 ACCCTTAGGAATTGAGTCACGTGTTGAGCCATTTCCTAGCTAGCAGTGCAGTTGG 6488472
*InMYB21B* KO #1 6488415 ACCCTTAGGAATTGAGTCACGTGTTGAGCCATTTCCTAGCTAGCAGTGCAGTTGG 6488469
*InMYB21B* KO #2 6487464 -----------------------------------CTAGCTAGCAGTGCAGTTGG 6487483
*InMYB21B* KO #3_1 6488350 ACCCTTAGGAATTGAGTCACGTGTTGAGCCATTTCCTAGCTAGCAGTGCAGTTGG 6488404
*InMYB21B* KO #3_2 6488407 ACCCTTAGGAATTGAGTCACGTGTTGAGCCATTTCCTAGCTAGCAGTGCAGTTGG 6488461
*InMYB21B* KO #4_1 6488408 ACCCTTAGGAATTGAGTCACGTGTTGAGCCATTTCCTAGCTAGCAGTGCAGTTGG 6488462

```
InMYB21B KO #4_2  6488416 ACCCTTAGGAATTGAGTCACGTGTTGAGCCATTTCCTAGCTAGCAGTGCAGTTGG 6488470
InMYB21B KO #5_1  6488391 ACCCTTAGGAATTGAGTCACGTGTTGAGCCATTTCCTAGCTAGCAGTGCAGTTGG 6488445
InMYB21B KO #5_2  6488416 ACCCTTAGGAATTGAGTCACGTGTTGAGCCATTTCCTAGCTAGCAGTGCAGTTGG 6488470
InMYB21B KO #6_1  6488391 ACCCTTAGGAATTGAGTCACGTGTTGAGCCATTTCCTAGCTAGCAGTGCAGTTGG 6488445
InMYB21B KO #6_2  6488414 ACCCTTAGGAATTGAGTCACGTGTTGAGCCATTTCCTAGCTAGCAGTGCAGTTGG 6488468

Wild type         6488473 AATCATGATGAAAAGGAAGCTCCAACTTGCTTTTG 6488505
InMYB21B KO #1    6488470 AATCATGATGAAAAGGAAGCTCCAACTTGCTTTTG 6488502
InMYB21B KO #2    6487484 AATCATGATGAAAAGGAAGCTCCAACTTGCTTTTG 6487517
InMYB21B KO #3_1  6488405 AATCATGATGAAAAGGAAGCTCCAACTTGCTTTTG 6488439
InMYB21B KO #3_2  6488462 AATCATGATGAAAAGGAAGCTCCAACTTGCTTTTG 6488496
InMYB21B KO #4_1  6488463 AATCATGATGAAAAGGAAGCTCCAACTTGCTTTTG 6488497
InMYB21B KO #4_2  6488471 AATCATGATGAAAAGGAAGCTCCAACTTGCTTTTG 6488503
InMYB21B KO #5_1  6488446 AATCATGATGAAAAGGAAGCTCCAACTTGCTTTTG 6488480
InMYB21B KO #5_2  6488471 AATCATGATGAAAAGGAAGCTCCAACTTGCTTTTG 6488503
InMYB21B KO #6_1  6488446 AATCATGATGAAAAGGAAGCTCCAACTTGCTTTTG 6488480
InMYB21B KO #6_2  6488469 AATCATGATGAAAAGGAAGCTCCAACTTGCTTTTG 6488501

Target sequence
PAM sequence
Replaced sequence
```

**Figure S7.** Alleles of *InMYB21B* in genome-edited lines

*InMYB21B* gene sequences in the wild-type and *InMYB21B* genome-edited lines. The guide RNA target sequences are highlighted in yellow, the PAM sequences are highlighted in blue, and replaced sequences are highlighted in gray.

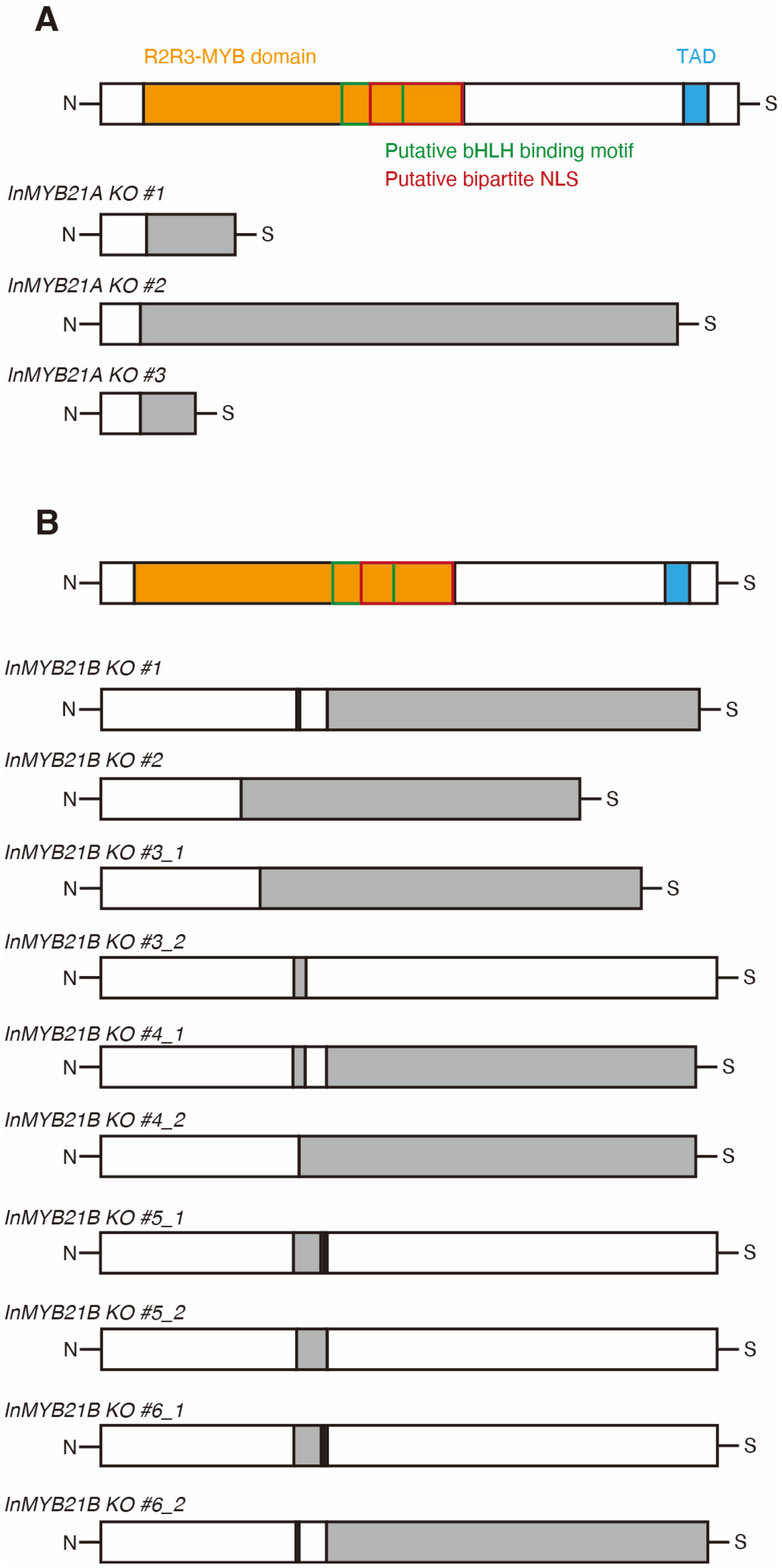
A
R2R3-MYB domain
TAD
N
S
Putative bHLH binding motif
Putative bipartite NLS
InMYB21A KO #1
InMYB21A KO #2
InMYB21A KO #3
B
InMYB21B KO #1
InMYB21B KO #2
InMYB21B KO #3_1
InMYB21B KO #3_2
InMYB21B KO #4_1
InMYB21B KO #4_2
InMYB21B KO #5_1
InMYB21B KO #5_2
InMYB21B KO #6_1
InMYB21B KO #6_2

**Figure S8.** Predicted amino acid sequences of genome-edited lines

Predicted R2R3-MYB domains (orange) and transcriptional activation domains (blue) of InMYB21A (**A**) and InMYB21B (**B**). Predicted amino acid sequences of *InMYB21A* and *InMYB21B* in genome-edited lines generated using CRISPR/Cas9. Amino acid sequences were predicted using ExPASy Translate. Regions with amino acid sequences identical to those of the wild type are shown in white, whereas regions with amino acid sequences different from those of the wild type are shown in gray.

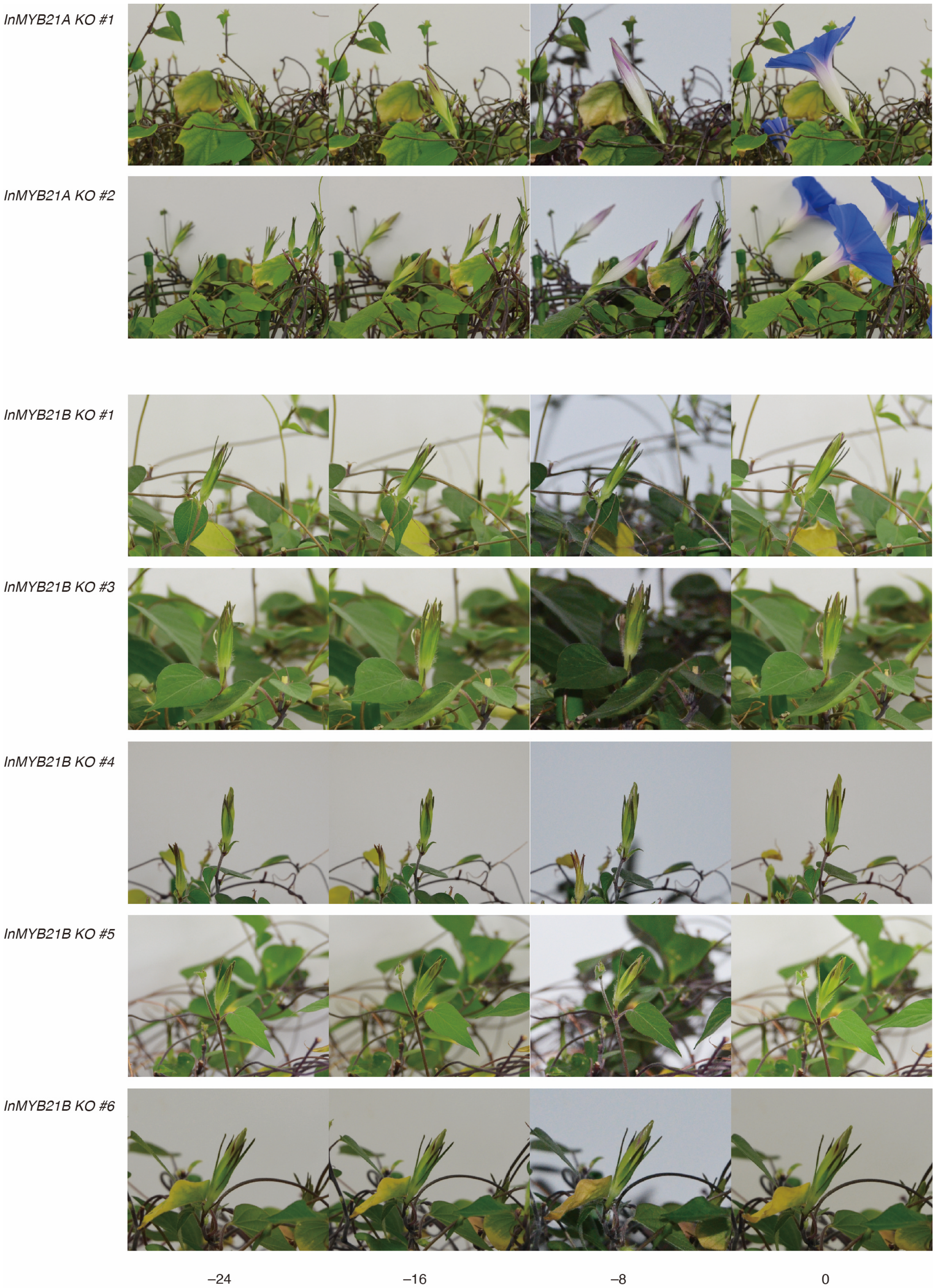


**Figure S9.** Time-lapse images of flower opening in *InMYB21A* KO and *InMYB21B* KO lines

Time-lapse images of *InMYB21A* KO lines #1 and #2 and *InMYB21B* KO lines #1 and #3–#6. Numbers below the images indicate the time relative to flower opening.

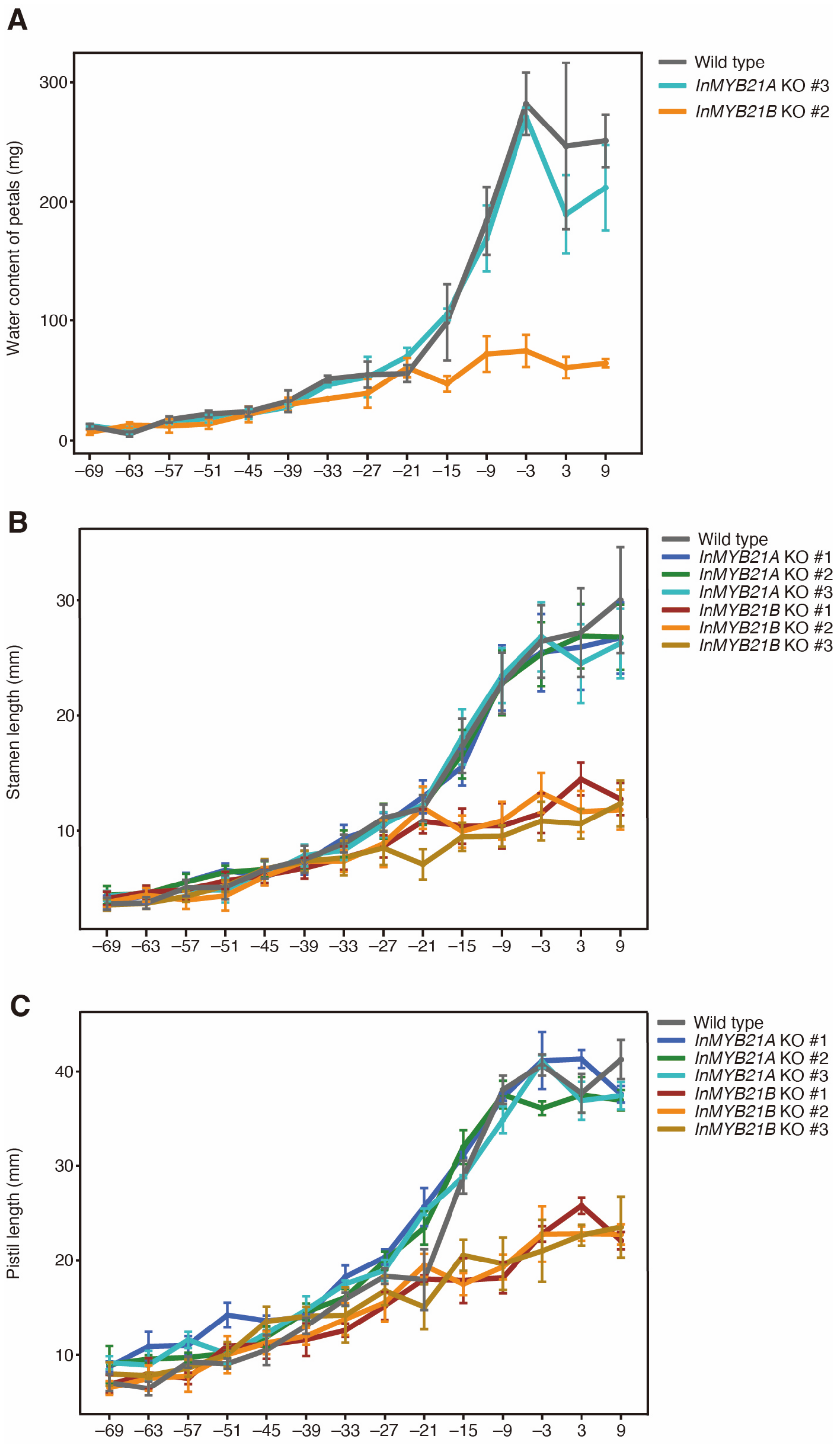


**Figure S10.** Temporal changes in petal water content, stamen length, and pistil length in wild-type, *InMYB21A* KO, and *InMYB21B* KO lines

Temporal changes in petal water content (**A**), stamen length (**B**), and pistil length (**C**). The x-axis indicates time relative to flower opening (0 h). Values are presented as means ± SD (n = 3) at each time point.

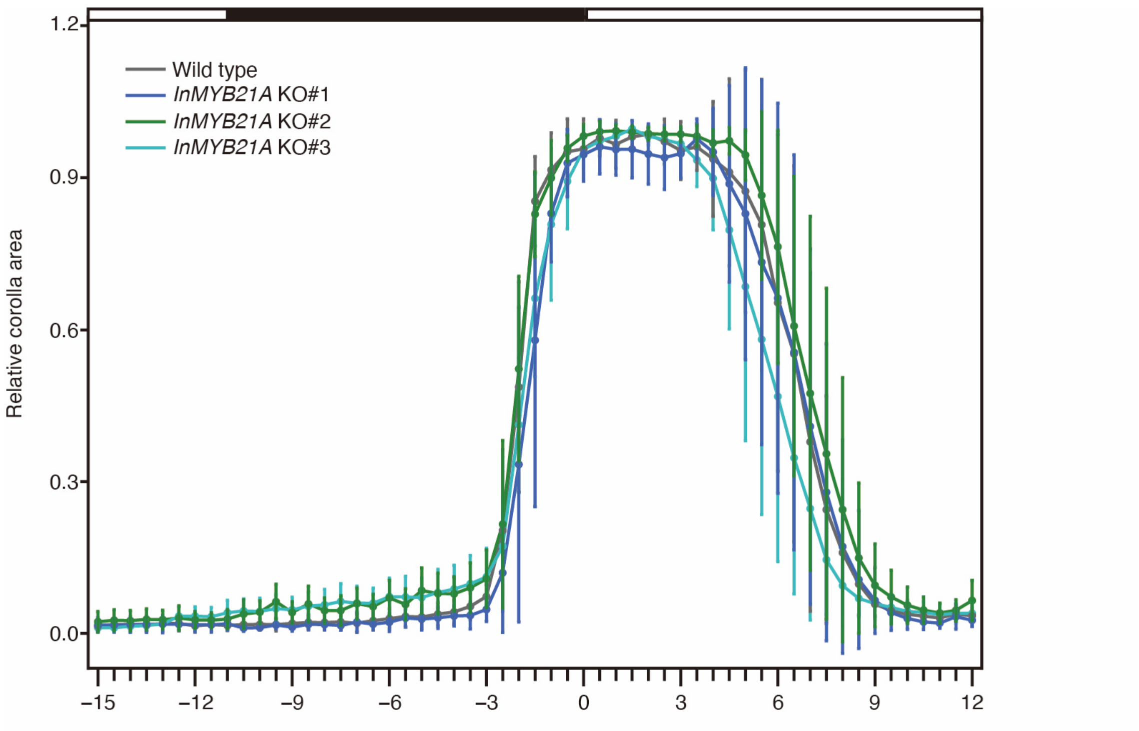


**Figure S11.** Measurement of relative corolla area

Temporal changes in relative corolla area in wild-type plants and *InMYB21A* KO lines #1–#3. The x-axis indicates time relative to flower opening (0 h), and the y-axis indicates the relative area of the pigmented region of the petals. Values are presented as means ± SE (n = 10). The bars at the top indicate light conditions, with white and black representing the light and dark periods, respectively. Relative corolla area was calculated by dividing the corolla area at each time point by the maximum corolla area of each flower.

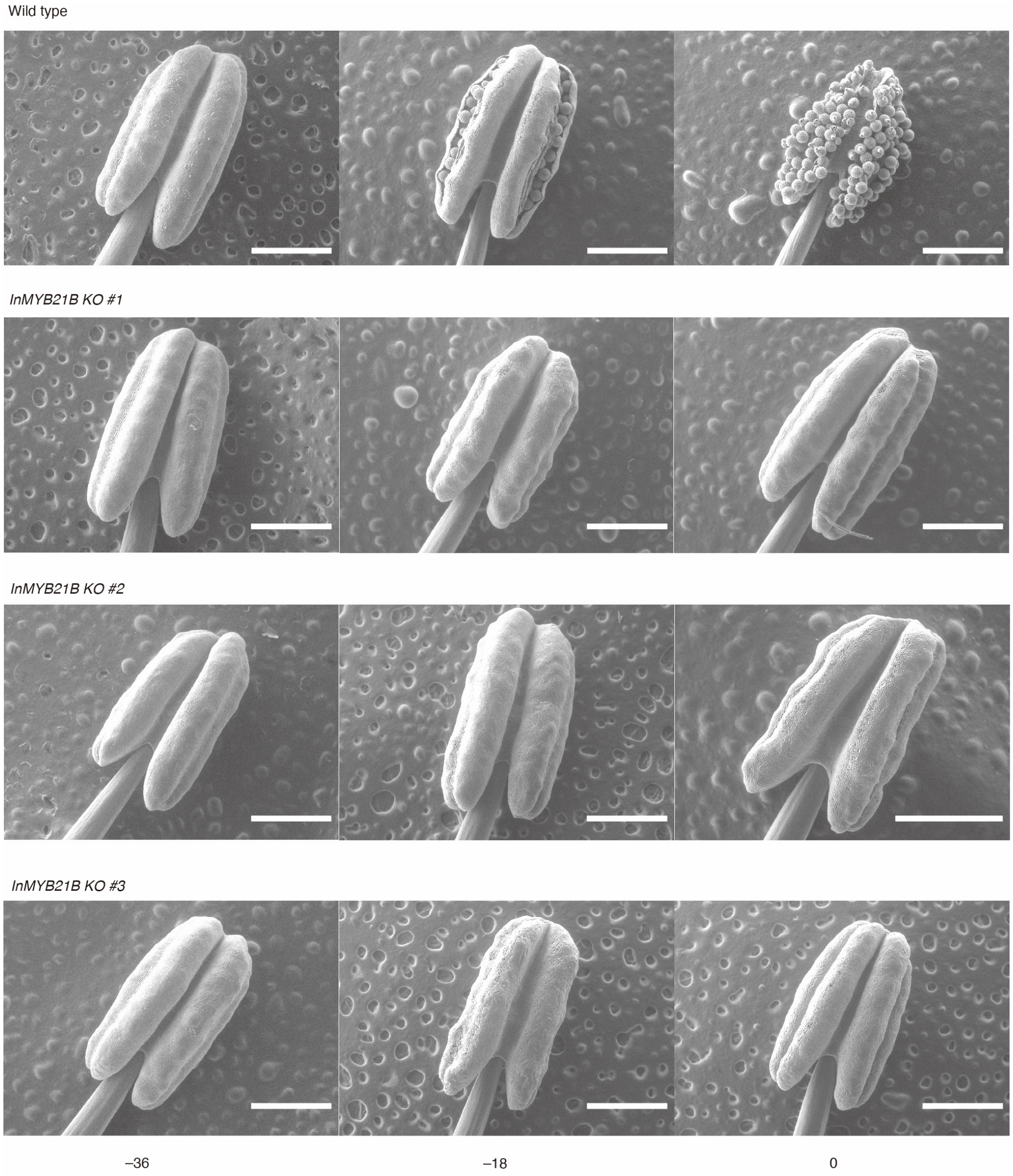


**Figure S12.** Scanning electron microscopy of anthers

Anthers of wild-type and *InMYB21B* KO lines were observed by scanning electron microscopy. The numbers below the images indicate the time relative to flower opening. Scale bars: 1 mm.

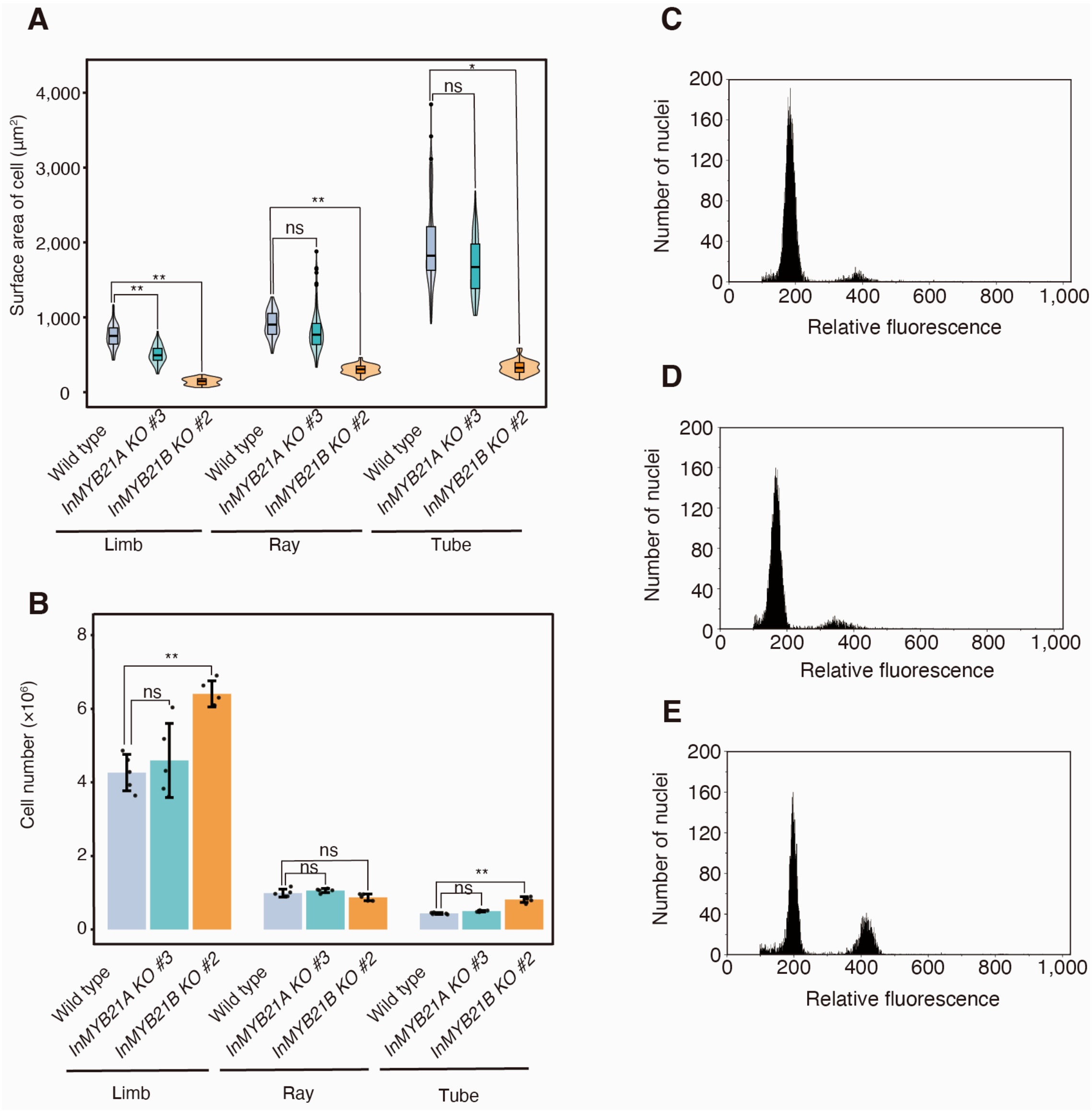


**Figure S13.** Measurement of epidermal cell size and cell number in petals and cell cycle analysis (**A**) Epidermal cell areas on the abaxial surfaces of the limb, ray, and tube in wild-type plants (blue), *InMYB21A* KO line #3 (green), and *InMYB21B* KO line #2 (orange), observed by scanning electron microscopy and quantified. Wild-type plants and each KO line were compared using Welch's t-test (n = 3; ns, not significant; $*p < 0.05$, $**p < 0.01$). (**B**) Epidermal cell numbers on the abaxial surfaces of the limb, ray, and tube in wild-type plants (blue), *InMYB21A* KO line #3 (green), and *InMYB21B* KO line #2 (orange). Wild-type plants and each KO line were compared using Welch's *t*-test (n = 5; ns, not significant; $*p < 0.05$, $**p < 0.01$). Flow cytometry histograms of wild-type plants (**C**), *InMYB21A* KO line #3 (**D**), and *InMYB21B* KO line #2 (**E**) at 3 h after flower opening.

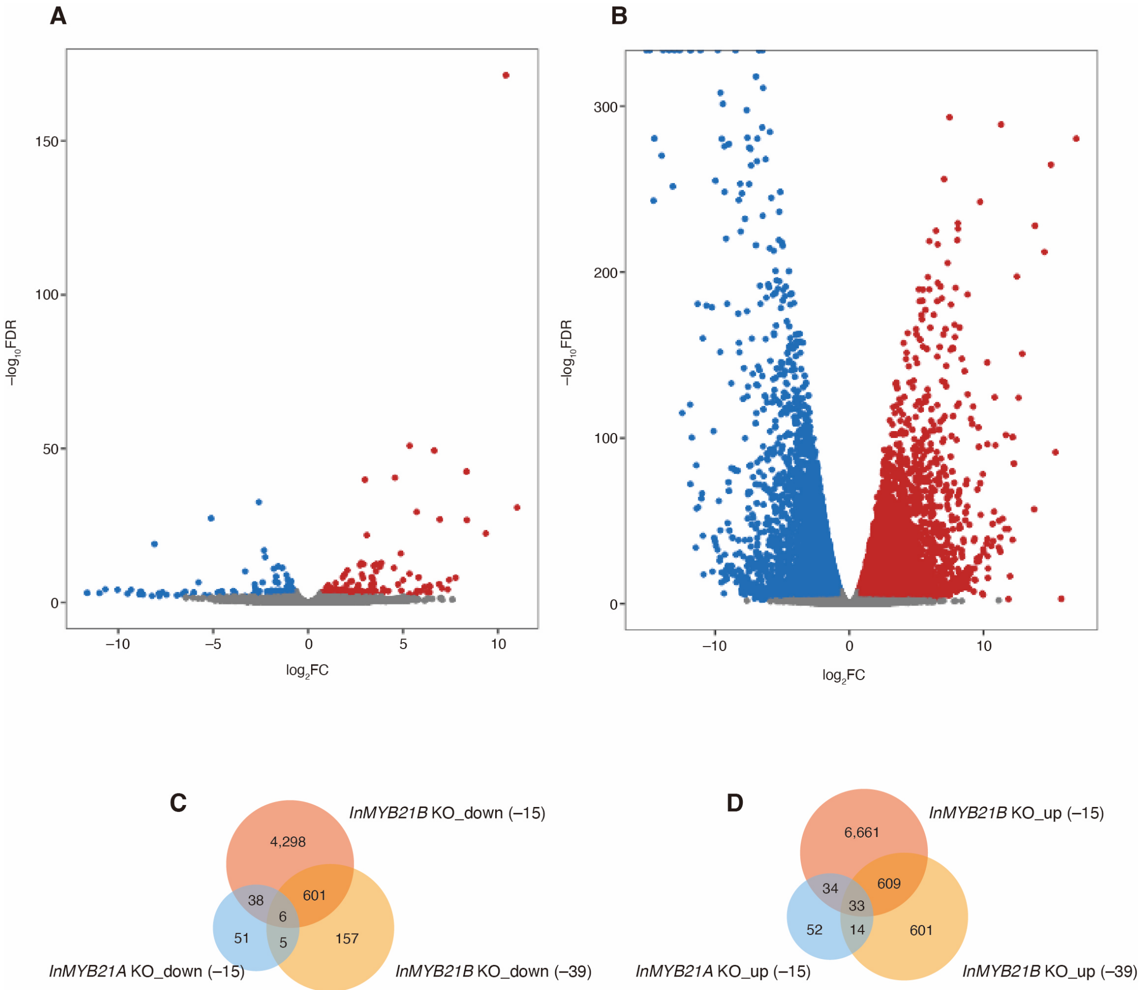


**Figure S14.** Comparative transcriptome analysis

(**A**) Volcano plot showing $\log_2$FC (x-axis) and $-\log_{10}$FDR (y-axis) for the comparative transcriptome analysis between *InMYB21A* KO line #3 and wild-type plants at 15 h before flower opening. Downregulated DEGs ($\log_2$FC < $-1$ and $-\log_{10}$FDR > 2) are shown in blue, upregulated DEGs (log2FC > 1 and $-$log10FDR > 2) in red, and other genes in gray. (**B**) Volcano plot showing $\log_2$FC (x-axis) and $-\log_{10}$FDR (y-axis) for the comparative transcriptome analysis between *InMYB21B* KO line #2 and wild-type plants at 15 h before flower opening. Downregulated DEGs ($\log_2$FC < $-1$ and $-\log_{10}$FDR > 2) are shown in blue, upregulated DEGs ($\log_2$FC > 1 and $-\log_{10}$FDR > 2) in red, and other genes in gray. (**C**) Venn diagram showing the overlap among downregulated DEGs in petals of *InMYB21A* KO line #3 at 15 h before flower opening and *InMYB21B* KO line #2 at 39 and 15 h before flower opening. (**D**) Venn diagram showing the overlap among upregulated DEGs in petals of *InMYB21A* KO line #3 at 15 h before flower opening and *InMYB21B* KO line #2 at 39 and 15 h before flower opening.

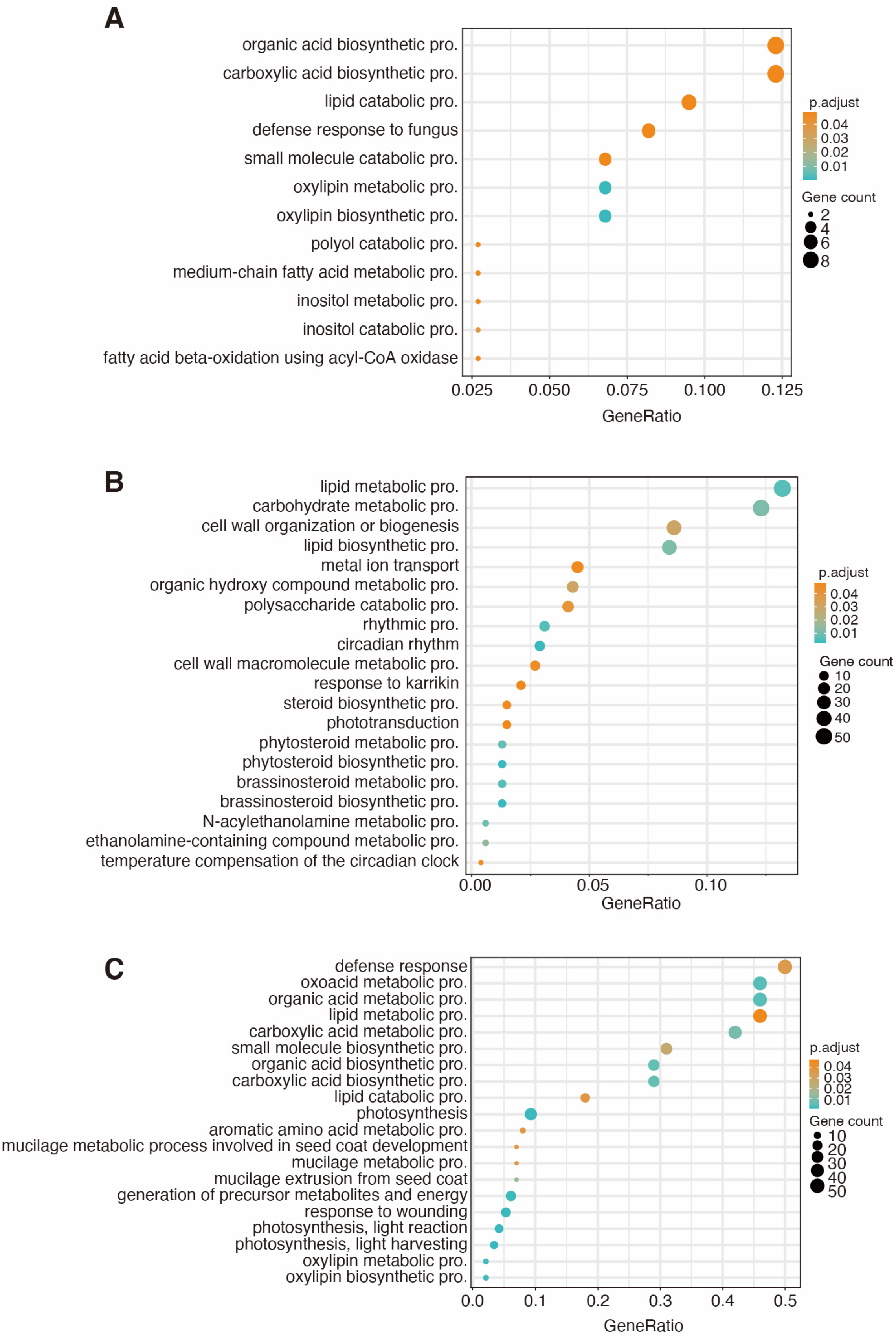
A
organic acid biosynthetic pro.
carboxylic acid biosynthetic pro.
lipid catabolic pro.
defense response to fungus
small molecule catabolic pro.
oxylipin metabolic pro.
oxylipin biosynthetic pro.
polyol catabolic pro.
medium-chain fatty acid metabolic pro.
inositol metabolic pro.
inositol catabolic pro.
fatty acid beta-oxidation using acyl-CoA oxidase
0.025
0.050
0.075
0.100
0.125
GeneRatio
p.adjust
0.04
0.03
0.02
0.01
Gene count
2
4
6
8
B
lipid metabolic pro.
carbohydrate metabolic pro.
cell wall organization or biogenesis
lipid biosynthetic pro.
metal ion transport
organic hydroxy compound metabolic pro.
polysaccharide catabolic pro.
rhythmic pro.
circadian rhythm
cell wall macromolecule metabolic pro.
response to karrikin
steroid biosynthetic pro.
phototransduction
phytosteroid metabolic pro.
phytosteroid biosynthetic pro.
brassinosteroid metabolic pro.
brassinosteroid biosynthetic pro.
N-acylethanolamine metabolic pro.
ethanolamine-containing compound metabolic pro.
temperature compensation of the circadian clock
0.00
0.05
0.10
GeneRatio
p.adjust
0.04
0.03
0.02
0.01
Gene count
10
20
30
40
50
C
defense response
oxoacid metabolic pro.
organic acid metabolic pro.
lipid metabolic pro.
carboxylic acid metabolic pro.
small molecule biosynthetic pro.
organic acid biosynthetic pro.
carboxylic acid biosynthetic pro.
lipid catabolic pro.
photosynthesis
aromatic amino acid metabolic pro.
mucilage metabolic process involved in seed coat development
mucilage metabolic pro.
mucilage extrusion from seed coat
generation of precursor metabolites and energy
response to wounding
photosynthesis, light reaction
photosynthesis, light harvesting
oxylipin metabolic pro.
oxylipin biosynthetic pro.
0.0
0.1
0.2
0.3
0.4
0.5
GeneRatio
p.adjust
0.04
0.03
0.02
0.01
Gene count
10
20
30
40
50

**Figure S15.** Gene Ontology (GO) enrichment analysis of differentially expressed genes (DEGs)

GO enrichment analysis of DEGs upregulated in *InMYB21A* KO line #3 at 15 h before flower opening (**A**), DEGs downregulated in *InMYB21B* KO line #2 at 39 h before flower opening (**B**), and DEGs commonly upregulated in *InMYB21B* KO line #2 at 39 and 15 h before flower opening (**C**). The significance threshold was set at corrected $p$-value < 0.05. GeneRatio indicates the proportion of genes in the analyzed DEG set assigned to each GO term. The size of each circle indicates the number of genes, and the color indicates the corrected $p$-value.

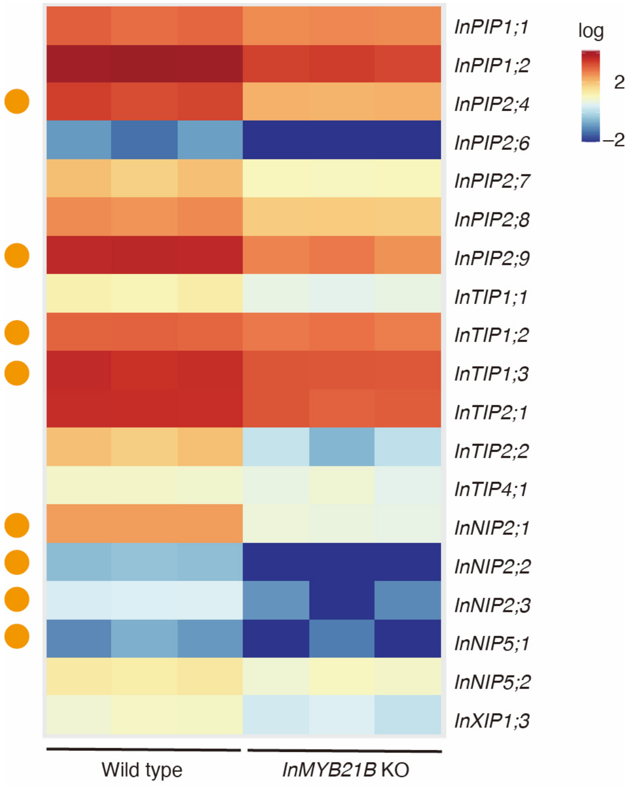


**Figure S16.** DEG analysis of aquaporin genes

Heatmap showing aquaporin genes downregulated in *InMYB21B* KO line #2 at 15 h before flower opening. Expression levels are shown as log10-transformed TPM values, with TPM values of 0 converted to 0.01 before transformation. Each column represents one biological replicate (n = 3). Genes marked with orange circles contain a cis-regulatory sequence of R2R3-MYB subgroup 19 within the 1,000-bp region upstream of the transcription start site.

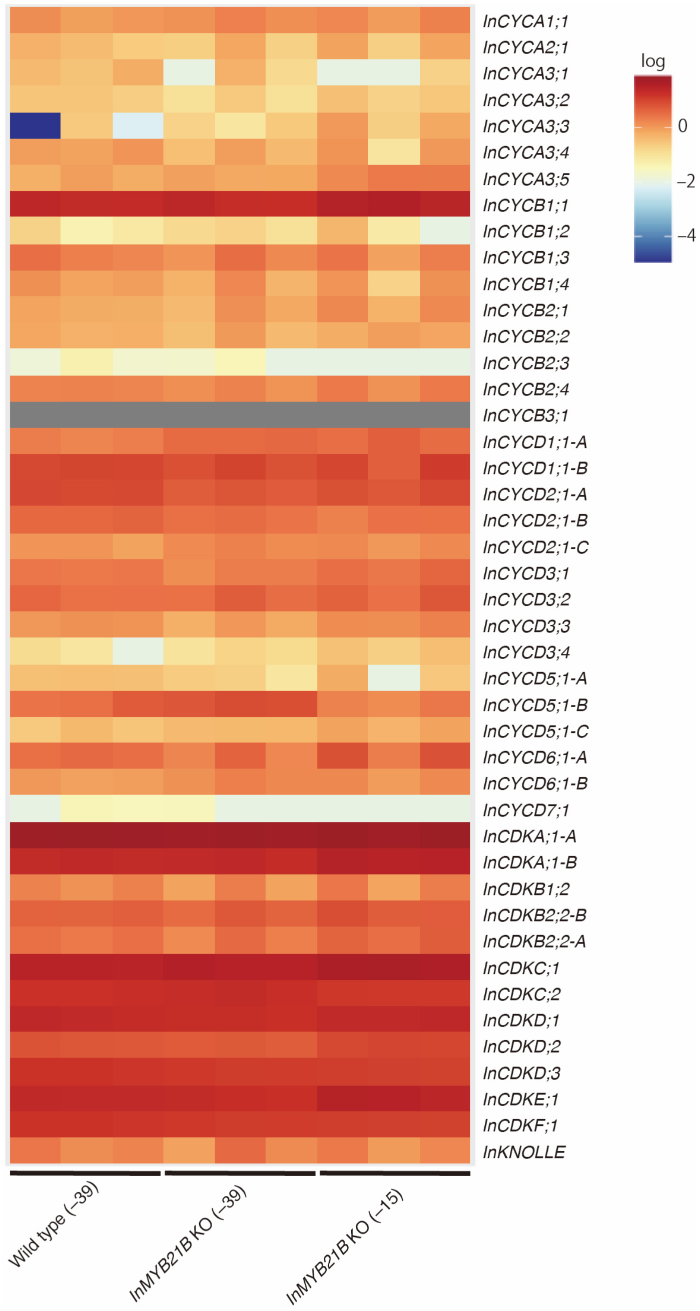


**Figure S17.** Heatmap analysis of cell cycle- and cell division-related genes

Heatmap showing the expression patterns of cell cycle- and cell division-related genes in wild-type plants at 39 h before flower opening, *InMYB21B* KO line #2 at 39 h before flower opening, and *InMYB21B* KO line #2 at 15 h before flower opening. Expression levels are shown as log10-transformed TPM values, with TPM values of 0 converted to 0.01 before transformation. Each column represents one biological replicate (n = 3).

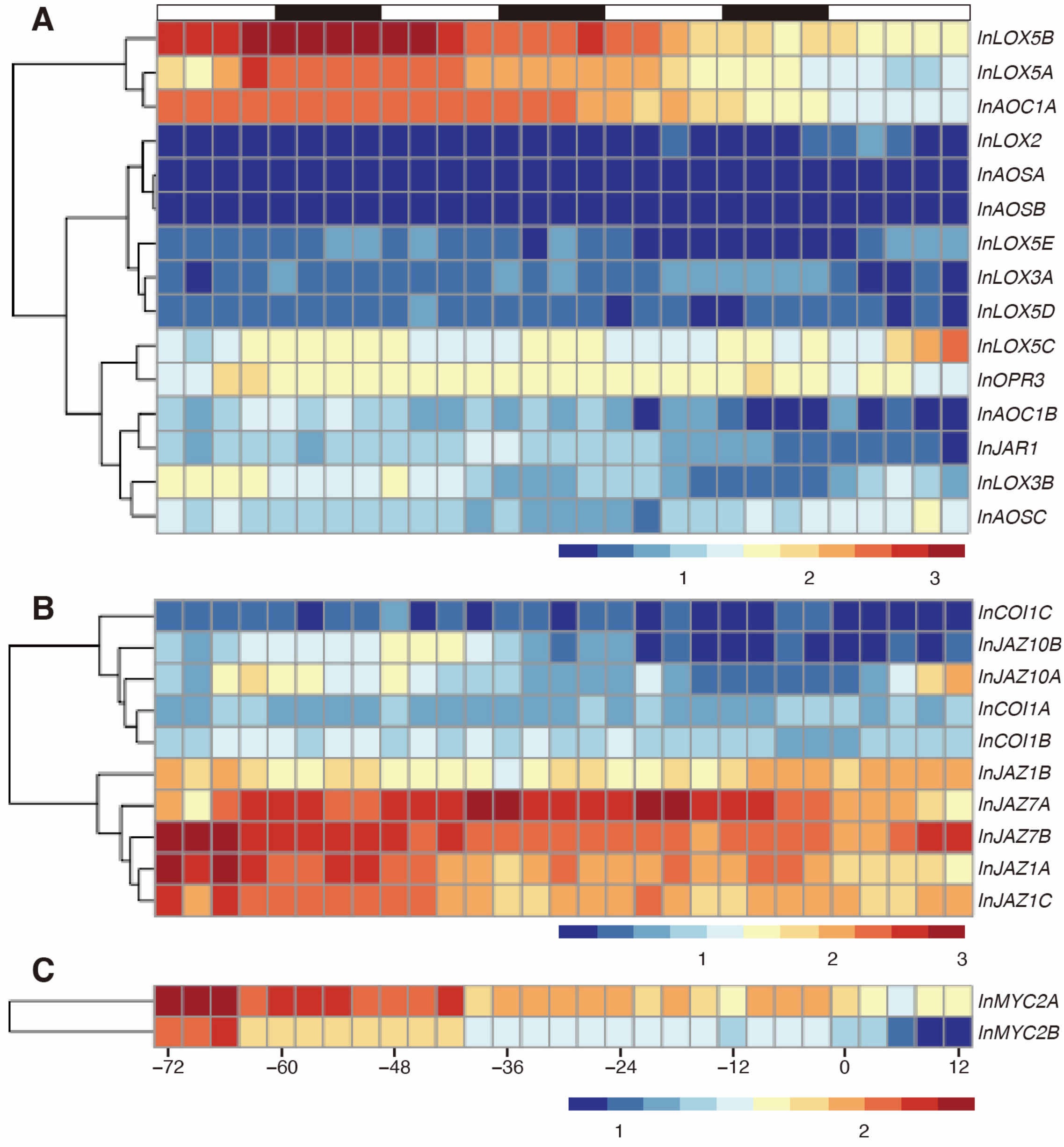


**Figure S18.** Heatmap analysis of the temporal expression patterns of jasmonate biosynthesis and signaling-related genes

Heatmaps showing the expression patterns of jasmonate biosynthesis genes (**A**), jasmonate signaling-related genes (**B**), and MYC genes (**C**). The gene lists are provided in Tables S20 and S21. Expression levels are shown as log10(FPKM + 1). The x-axis indicates time relative to flower opening (0 h). The bars at the top indicate light conditions, with white and black representing light and dark periods, respectively.
